# A Non-Viral CRISPR/Cas9 HDR Platform for Stable Engineering of Solid Tumor Models

**DOI:** 10.64898/2026.06.01.729035

**Authors:** Shoaib Afzal, Maximilian Pilgram, Julia Macos, Emily Ohlendorf, Lena Raab, Jonas Kath, Viktor Glaser, Ana-Maria Nitulescu, Casper F T van der Ven, Chahrazad Lachiheb, Maria Stecklum, Norman Michael Drzeniek, Kathleen Anders, Dimitrios Laurin Wagner, Ralf Kühn, Annette Künkele, Michael Launspach

## Abstract

Virus-free genome engineering provides a flexible alternative to viral vectors for generating genetically modified cell models. Here, we establish an integrated biosafety level 1-compatible CRISPR/Cas9 homology-directed repair (HDR) workflow for stable transgene knock-in in neuroblastoma cell lines using non-viral delivery approaches. We systematically evaluated donor cassette architecture and delivery conditions across electroporation-based Cas9 ribonucleoprotein (RNP) delivery and lipid nanoparticle (LNP)-mediated co-delivery of Cas9 mRNA, sgRNA, and donor DNA. Modular AAVS1-targeting donor constructs identified a compact EF1α(s)-Donor-Q8-Tag-sPA cassette that consistently yielded the strongest HDR-associated knock-in readouts, achieving up to 60% stable reporter-positive cells following electroporation without HDR enhancers. While LNP-mediated delivery enabled efficient CRISPR cargo co-delivery and generation of genetically modified tumor cell populations, knock-in efficiencies remained lower than those observed with electroporation. Subsequent enrichment approaches enabled generation of highly pure edited cell populations following both delivery strategies. Functional validation demonstrated stable transgene expression in vitro, including in three-dimensional bioprinted tumor models, and in vivo in xenograft mice without impairing tumor growth or viability. Together, these findings establish a practical non-viral HDR platform for stable engineering of solid tumor models and provide a framework for further optimization of genome editing workflows across distinct delivery modalities.

**Key findings:**

- We establish a complete, virus-free CRISPR/Cas9 HDR workflow that reliably enables stable knock-in in solid tumor cell lines, demonstrated here in two neuroblastoma models under biosafety level 1 conditions.
- We establish and evaluate LNP-mediated co-delivery of Cas9 mRNA, gRNA, and donor DNA for non-viral HDR knock-in in solid tumor models, revealing delivery modality-specific differences in editing efficiency, toxicity, and expression dynamics.
- By systematically varying donor architectures, we identify a compact HDR template - combining a shortened custom EF1α promoter, the minimal Q8 surface reporter, and a synthetic polyadenylation signal (sPA) - that markedly improves knock-in efficiency in solid tumor cell lines, outperforming conventional cassettes.
- Virus-free edited tumor cells generated using this workflow retain stable transgene expression and functional fitness in 3D bioprinted tumor constructs and xenograft mouse models, directly linking in vitro knock-in optimization to in vivo relevance.
- The resulting biosafety level 1 compatible, end-to-end pipeline - integrating donor design, digital PCR-based quantification of precise integration, and enrichment strategies-offers a practical and transferable platform for engineering transgenic solid tumor models without viral vectors.

## Introduction

Genetic engineering has profoundly advanced the development of cellular therapies, particularly in cancer immunotherapy, with targeted “knock-in” approaches that insert exogenous DNA sequences into specific genomic loci proving crucial for both disease modeling and therapeutic innovation^1^. Transgenic and isogenic cell lines produced through precise genome editing enable detailed functional studies of oncogenic pathways, systematic identification of therapeutic vulnerabilities, and high-fidelity screening of anti-cancer compounds^2^. Historically, the generation of such cell lines has relied heavily on viral and transposon-based delivery systems. Lentiviral vectors, derived from HIV-1, offer high transduction efficiency and stable integration even in non-dividing cells^3^. Yet their retroviral origins raise serious biosafety concerns, including risks of insertional mutagenesis and disruption of essential genes^4^. Adeno-associated virus (AAV) vectors offer a safer profile due to largely episomal persistence and low immunogenicity but suffer from a small packaging limit of around 4.7 kb, constraining the complexity of therapeutic cassettes^5^. Nonviral systems like Sleeping Beauty or PiggyBac transposons circumvent viral risks but still integrate semi-randomly and require transposase activity, raising the potential for genomic instability^6^. Even after decades of refinement, viral-vector gene therapies continue to contend with treatment-related adverse events and genotoxicity risks, underscoring the demand for next-generation platforms that combine high efficiency with precise, site-specific integration^7^.

CRISPR-Cas9-mediated homology-directed repair (HDR) has emerged as a transformative technology for introducing precise genomic modifications using exogenous donor templates^8^. Unlike viral vectors, CRISPR enables targeted insertion with minimal genomic footprint, reducing risks of insertional mutagenesis and allowing predictable expression of integrated transgenes^9,10^. However, HDR efficiency remains a major bottleneck, with rates often below 10-15% in bulk populations without selection, particularly in non-dividing or postmitotic cells^11^. Advances in donor design, such as end-modifications, optimized homology arms, asymmetric single-stranded templates, or covalent tethering of donor DNA to Cas9 RNPs have yielded increases in HDR efficiency^12–15^. Importantly, optimal edits are achieved when the desired modification is close to the Cas9-induced cut site, a principle supported by microhomology analyses and systematic studies on edit-to-cut distances^11,16^. Furthermore, optimizing the delivery complex itself is critical; the use of anionic polymers like polyglutamic acid (PGA) to form stabilized complexes of Cas9 RNP and donor DNA has been shown to be a highly effective strategy, markedly improving knock-in efficiency while simultaneously increasing the viability of sensitive primary cells after electroporation^2,17^. In cancer immunotherapy, these precise genome-editing tools are increasingly vital for addressing challenges in adoptive cell therapies like CAR T cells, which face hurdles such as insufficient tumor infiltration, T cell exhaustion, and antigen escape^18,19^. Beyond simply integrating CAR constructs, CRISPR knock-ins can enable programmable secretion of cytokines or chemokines, improving T-cell trafficking and function^20^. Reviews emphasize the need for multi-antigen targeting, armored CAR designs, and localized cytokine release to overcome the immunosuppressive tumor microenvironment^21^. Virus-free generation of engineered immune cells, such as TRAC-locus replacement with Cas9 RNPs and DNA donors, is a key innovation that enhances safety by avoiding viral integrations^2^. Similarly, tumor cell lines engineered to express chemokines, costimulatory molecules or neoantigens may offer new platforms for studying and overcoming barriers in solid tumor immunotherapy and preclinical disease modeling. Beyond HDR-mediated knock-ins, emerging technologies such as base editing, prime editing, and RNA-guided integrases promise precise genome modifications without inducing double-strand breaks^22^. Prime editing, for example, enables targeted insertions and small sequence changes without double-strand breaks, instead relying on a nick-and-repair process, although efficiency for large knock-ins remains limited^22,23^. Similarly, CRISPR-associated transposase systems offer programmable DNA integration but are still in early development for therapeutic applications. While these tools hold significant future promise, robust protocols for large, precise insertions in complex cell systems like tumor lines or immune cells are not yet established, underscoring the continued importance of optimizing HDR-based approaches for clinical translation^24^.

Lipid nanoparticles (LNPs) have recently emerged as a powerful alternative to electroporation for delivering CRISPR reagents^25,26^. Initially propelled into prominence by mRNA vaccines, LNPs now show promise for delivering Cas9 mRNA, RNPs, and DNA donors without genome integration risks associated with viral vectors^27^. Clinical proof-of-concept was demonstrated in transthyretin amyloidosis patients treated with NTLA-2001, an LNP-based in vivo CRISPR therapy^28^. More recently, LNPs have been used for in vivo CAR T-cell generation in preclinical models^29^. Despite progress, LNP co-delivery of Cas9 RNPs and large DNA donors remains challenging, especially for achieving high-efficiency HDR in solid-tumor models or primary immune cells^27,29^. Knock-in rates remain low, necessitating labor-intensive screening; most studies optimize single variables without exploring their interdependence; and long-term genomic stability and off-target risks remain incompletely characterized^30,31^. Comprehensive frameworks integrating functional assays, genomic analyses, and delivery optimization are urgently needed. In this work, we address these challenges by systematically comparing promoter-transgene cassettes of varying sizes across two biologically distinct neuroblastoma lines using RNP electroporation platforms and optimized LNP formulations for Cas9 and donor co-delivery; quantifying on-target and off-target integrations using digital PCR; and validating functional transgene expression both in 3D models and in vivo. Our integrated approach aims to boost knock-in efficiency, ensure genomic integrity, and establish quality-control pipelines and state-of-the-art methodical protocols essential for preclinical as well as translational applications.

## Material & Methods

### Cell Culture

SK-N-BE(2)-C (ATCC CRL-2268) and SK-N-AS (ATCC CRL-2137) neuroblastoma cell lines were kindly provided in 2018 by PD Dr. med. Hedwig Deubzer (Charité – Universitätsmedizin Berlin) and Prof. Michael Claus V. Jensen (Seattle Children’s Hospital), respectively. SK-N-AS cells were maintained in RPMI 1640 medium (Gibco), while HEK293T (ATCC CRL-3216) and SK-N-BE(2)-C cells were cultured in high-glucose DMEM (Gibco). All cultures were kept at 37°C in a humidified atmosphere with 5% CO₂ and supplemented with 10% heat-inactivated fetal bovine serum (FBS). All cell lines were routinely monitored for bacterial, fungal, and mycoplasma contamination using standard microbiological screening protocols. Furthermore, the neuroblastoma cell lines were periodically authenticated by short tandem repeat (STR) profiling to confirm their tumor identity, with the most recent authentication performed in 2024. After knock-in experiments, media included 1% penicillin/streptomycin (Gibco). For LNP application, cell line media was replaced with ImmunoCult TM-XF T Cell expansion media including 1% penicillin/streptomycin (Gibco). To enable LDL receptor-mediated endocytosis, 1% ApoE3/4 was added.^32^

### Cloning and HDR Template Production

Transgene constructs (Supplementary Table 1) were designed in SnapGene to optimize cassette architecture, including promoters, coding sequences, regulatory elements, and polyadenylation signals, with codon optimization and removal of cryptic splice sites (Supplementary Table 2). Overlapping primers (∼15 bp homology arms) were designed using SnapGene and Primer-BLAST, aiming for Tm 58–62°C and minimal secondary structures (Supplementary Table 3). Constructs were assembled via NEBuilder HiFi DNA Assembly (NEB) without restriction digestion for scarless assembly. PCR amplifications used KAPA HiFi Polymerase (Roche); products were gel-verified, purified with QIAquick Gel Extraction Kit, and quantified. Linearized blunt-end plasmid backbones (Twist Bioscience) and PCR fragments were combined with 2× NEBuilder HiFi Master Mix at a 1:2 vector-to-insert molar ratio (50 ng vector) in 10 μl reactions, incubated at 50°C for 1 h. XL10-Gold E. coli were transformed with 2 μl assembly mix, plated on LB-ampicillin agar, and incubated overnight at 37°C. Colony PCR screened clones for correct inserts, followed by plasmid purification (ZymoPURE Miniprep Kit, Zymo Research). Constructs were verified by Sanger sequencing (LGC Genomics) to confirm homology junctions, promoters, and coding regions. For HDR template generation, verified plasmids served as PCR templates in 400 µl reactions to amplify dsDNA fragments flanked by ∼400 bp homology arms using KAPA HiFi polymerase^33^. Products were purified with AMPure XP beads (Beckman Coulter) for high-purity genome editing applications. In addition, ssDNA HDR templates were used for comparison with dsDNA, produced as described in the Supplementary Methods and according to previously published protocols^34^.

### mRNA In Vitro Transcription and Handling

For linear dsDNA, touchdown PCR was optimized with KAPA HiFi HotStart ReadyMix to produce EGFP and CRISPR/Cas9_mCherry templates^35^. In vitro transcription used HiScribe T7 High-Yield Kit (NEB), with full N1-methyl-pseudouridine substitution and cotranscriptional capping (CleanCap AG, TriLink). Residual DNA was removed with TURBO DNase (Invitrogen), and RNA purified using Monarch RNA Cleanup Kit. RNA was quantified via NanoDrop and Qubit RNA BR assays.

### Electroporation for CRISPR/Cas-Mediated Transgene Knock-In

Cells were checked for morphology and ∼80% confluency. Media were pre-warmed, and either the 4D Nucleofector X Unit (Lonza) or Neon Transfection System (Thermo Fisher) prepared per manufacturer protocols. RNP complexes were freshly assembled by combining an aqueous solution of 15-50 kDa poly-L-glutamic acid (100µg/µl) and gRNA (3,125µg/µl (1µg = 32pmol)), then adding SpCas9 (10 µg/µl (1µg = 6pmol)). Typical molar ratios were 2:1 gRNA:Cas9 (e.g., 48 pmol:24 pmol for Lonza; 15-24 pmol gRNA with 5-12 pmol Cas9 for Neon) as used by Nguyen et al. (2020)^17^. For knock-in experiments using the Lonza system in a 20 µl volume, 50 µg of PGA was used per reaction. This resulted in PGA:gRNA mass ratios of 33:1 for SK-N-BE(2)C cells (0.6 × 10⁶ per reaction; 1.5 µg gRNA) and 25:1 for SK-N-AS cells (0.8 × 10⁶ per reaction; 2.0 µg gRNA). For the Neon system, reactions were prepared in a 12 µl total volume for 0.5 × 10⁶ cells. Each reaction contained 25 µg PGA and 0.75 µg gRNA, maintaining a consistent PGA:gRNA mass ratio of 33:1 across both cell lines. Mixtures were incubated at room temperature for 15–30 min. HDR templates (1.2–2 μg per 10⁶ cells) were added, gently mixed, and kept chilled. Lonza 4D Nucleofection: Cells were trypsinized, washed twice with PBS, and resuspended in SF buffer. For each reaction, 20 μl cell suspension was mixed with RNP-HDR DNA complex, loaded into Nucleocuvette Strips, and electroporated using cell-specific programs optimized for each cell type (SK-N-AS FF104, SK-N-BE(2)-C DN110). Post-nucleofection, 150 μl RPMI-1640 with 10% FCS was added, avoiding DMEM due to potential calcium toxicity. After 10 minutes at 37°C, cells were transferred to pre-warmed culture plates. Media were refreshed at 24 h, particularly if HDR Enhancer (IDT) was used. Neon Transfection: Cells were dissociated with 0.05% trypsin-EDTA, washed in PBS, and resuspended in Buffer R (6 μl per reaction). Neon transfections were prepared with Buffer E and cleaned Neon Tips. Cell suspensions (5 μl) were mixed with 7 μl RNP–HDR mix, aspirated into Neon Tips without air bubbles, and electroporated using optimized parameters. Post-electroporation, cells were transferred into pre-warmed RPMI-1640 with 10% FCS and antibiotics (e.g., 100 U/mL penicillin, 100 μg/mL streptomycin). For both systems, cell viability and morphology were assessed at 24 h post-nucleofection, and cultures transitioned to complete medium with antibiotics.

### LNP Formulation and Characterization

LNPs were formulated on the NanoAssemblr Spark platform (Precision Nanosystems) under sterile conditions after purging the instrument. Lipid mixes (both manufacturer kits and custom ALC-0315 (ALC) formulations) were thawed at 55°C for 5 min and equilibrated to room temperature (Supplementary Table 5). All nucleic acids (HDR template DNA, mRNA, and gRNA) were prepared at 1 μg/μl concentration. For knock-in experiments, nucleic acid mixes contained 2.40 μl HDRT DNA, 1.80 μl gRNA, 2.40 μl mRNA, 2.11 μl Formulation Buffer I (for kits) or 100 mM Na-acetate pH 4 (for custom formulations), and 12.30 μl RNase-free water. LNPs were formulated by loading the Spark cartridge with Dilution Buffer/PBS, nucleic acid mix and lipid mix. Standard formulations used 28.8 μl Dilution Buffer/PBS, 19.2 μl nucleic acid mix and 9.6 μl lipid mix, for formulation device setting 2 was used. Post-assembly, LNPs were diluted in 48 μl Dilution Buffer/PBS. For custom lipid mixes, individual lipids were dissolved in 100% ethanol in a bench top shaker at 40°C (Cholesterol 25µg/µl, 18:DSPC 25µg/µl, DMG-PEG(2000) 25µg/µl, DSPE-PEG(2000) Maleimide 10µg/µl, ALC-0315 (25µg/µl)). Desired volumes of lipid stock solutions were combined in DNAse/RNAse-free vials, mixed, and stored at –80°C. Multiple custom ALC-based formulations were generated to evaluate whether non-proprietary LNP compositions with differing physicochemical properties could support CRISPR cargo delivery and HDR-associated editing outcomes in neuroblastoma cells: LMA (own formulation), LMB based on Rosenblum et al. 2020^36^, and LMC based on Kenyo et al. 2021^37^. LMB contains DSPE-PEG(2000)-maleimide to enable potential future conjugation of thiol-containing targeting ligands (e.g., monoclonal antibodies or peptides). In the experiments reported here, no thiol-bearing molecules were added, and the maleimide functionality was not exploited. The low reaction probability under the experimental conditions and short incubation times makes nonspecific maleimide–thiol interactions with media components or cell-surface proteins unlikely. Each formulation was prepared at four molarities (35 mM, 25 mM, 16 mM, and 10 mM) under identical NanoAssemblr conditions. No post-formulation dialysis was performed. Encapsulation efficacy was determined using the Quant-iT™ RiboGreen RNA Assay Kit (Invitrogen™) following the manufacturer protocol. LNP size, polydispersity index (PDI), peak 1 area, were measured by dynamic light scattering (DLS) on the Zetasizer Ultra (Malvern Panalytical), with samples diluted 1:200 in nuclease-free HPLC-grade water.

### 3D Bioprinted Tumor Infiltration Model

Following CRISPR/Cas nucleofection and establishment of stable transgene-expressing cell lines, we generated standardized three-dimensional neuroblastoma constructs through collaboration with Cellbricks GmbH (Berlin, Germany)^38^. Using a hydrogel-based bioink optimized for neuroblastoma culture, SK-N-AS and SK-N-BE(2)-C cells - either unmodified or engineered to secrete immunostimulatory cytokines - were encapsulated in cylindrical scaffolds (3 mm diameter × 1 mm height; 7.07 mm³ volume) to ensure uniform cell distribution and reproducible tumor architecture.

### Housing and Handling of Animals and Xenograft Mouse Model

All animal studies followed German Animal Welfare regulations and were approved by LAGeSo Berlin (approval: Anz.Ther.: Reg E0023-23). Mice were kept under a 12-hour light/dark cycle at 23°C with ad libitum food and water. Animals were monitored twice daily. For xenografts, 5 × 10⁶ tumor cells in Matrigel were injected subcutaneously into 6–8-week-old female NOD.Cg-Prkdcscid Il2rγtm1Sug/JicTac mice (CIEA NOG; Taconic Biosciences, Inc.). Tumor growth was tracked biweekly by caliper measurements, and volume calculated as *V* = (length × width²)/2. Blood was collected via retrobulbar plexus, processed, and serum stored at –80°C. Tumors were excised, weighed, and split for formalin-fixation or snap-freezing.

### Statistical Analysis

Statistical analyses were performed in GraphPad Prism v10.2.0. Data are shown as mean ± SD unless noted. Comparisons among multiple groups used One-Way ANOVA or Kruskal-Wallis tests with Dunn’s correction; two-group comparisons used Mann-Whitney tests. Time-course analyses employed two-way ANOVA with Tukey’s post hoc tests. Correlations used linear regression and Spearman’s rank tests. Growth curves were fitted to logistic models. For dPCR quantification, two-way ANOVA was performed. Significance was set at p < 0.05. Exact sample sizes are detailed in figure legends.

Additional methods are described in the supplementary methods section. Equipment, consumables, antibodies and software used are listed in Supplementary tables 6-8.

## Results

### Modular Vector Testing Reveals Cell Line-Dependent Transgene Expression and Confirms Smaller Custom Components Are as Effective as Standard Designs

We employed a CRISPR/Cas9 homology-directed repair (HDR) workflow targeting the AAVS1 safe harbor locus to enable virus-free, stable transgene knock-in in neuroblastoma cells (Figure 1a)^33^. This system integrates Cas9 and gRNA with donor DNA templates flanked by 400 bp homology arms, supporting precise transgene insertion and modular construct design (Figure 1b). (Supplementary Figure S1a-c). Phase-contrast microscopy revealed morphological differences between cell lines, with SK-N-AS exhibiting a fibroblast-like, elongated shape and SK-N-BE(2)-C displaying a more epithelial, clustered morphology. These intrinsic morphological differences influence membrane area, cytoskeletal organization, and uptake dynamics, which likely contribute to the lower transfection efficiency and reduced HDR performance observed in SK-N-AS. (Supplementary Figure S1d). Before CRISPR knock-in experiments, we systematically evaluated modular donor components in transient transfection assays using Effectene and flow cytometry (Supplementary Figure S2a). Testing four promoters (CMV, MND, EF1α, and a custom designed shortened EF1α(s)) showed that in SK-N-AS cells, all promoters produced modest transgene expression, with 10–20% positive cells and similar mean fluorescence intensities (MFI). Conversely, SK-N-BE(2)-C cells consistently achieved higher expression across all promoters, reaching 60–100% positive cells, independent of promoter choice (Figure 1c). Statistical analysis confirmed that manually shortening the EF1α promoter did not significantly affect expression within each cell line (p > 0.05) (Figure 1c). Reporter benchmarking showed that the compact Q8 surface epitope tag^39^ performed comparably to standard GFP and RFP reporters - which exhibited cell-line–specific variation in expression and detectability - indicating that Q8 was a suitable and robust candidate for downstream knock-in experiments (Figure 1d and Supplementary Note 1). Polyadenylation signal choice had no significant impact, as constructs with bovine growth hormone (bGH) or synthetic polyadenylation signals (sPA) performed comparably in both cell lines which allowed for choosing the smaller sPA signal for downstream applications (Supplementary Figure S2c). While E2A and P2A self-cleaving-peptides showed no overall differences, reporter positioning within bicistronic constructs proved crucial. In SK-N-BE(2)-C, but not SK-N-AS, placing the reporter second downstream of CXCL10-P2A reduced expression, suggesting translational inefficiency or incomplete peptide cleavage (Supplementary Figure S3a, b). Western blot confirmed incomplete cleavage, and ELISA revealed lower CXCL10 secretion when CXCL10 was downstream of the reporter. Incomplete 2A-mediated peptide cleavage likely reduces translation of downstream elements which highlights the importance of gene order in bicistronic cassettes, particularly in solid tumor cells with lower global translation efficiency. (Supplementary Figure S3c, d). Functional validation demonstrated robust CXCL10 secretion and bioactivity, with ELISA detecting high chemokine levels and transwell assays confirming T-cell chemotaxis induced by secreted CXCL10 (Supplementary Figure S3d, e). These results show that while promoter choice minimally affects expression within a cell line, transgene expression efficiency depends strongly on cell-intrinsic factors and vector architecture. Notably, smaller custom elements - including EF1α(s), sPA, and the Q8 reporter - performed comparably to standard components, supporting their suitability for modular, virus-free genome editing applications.

**Figure 1.**
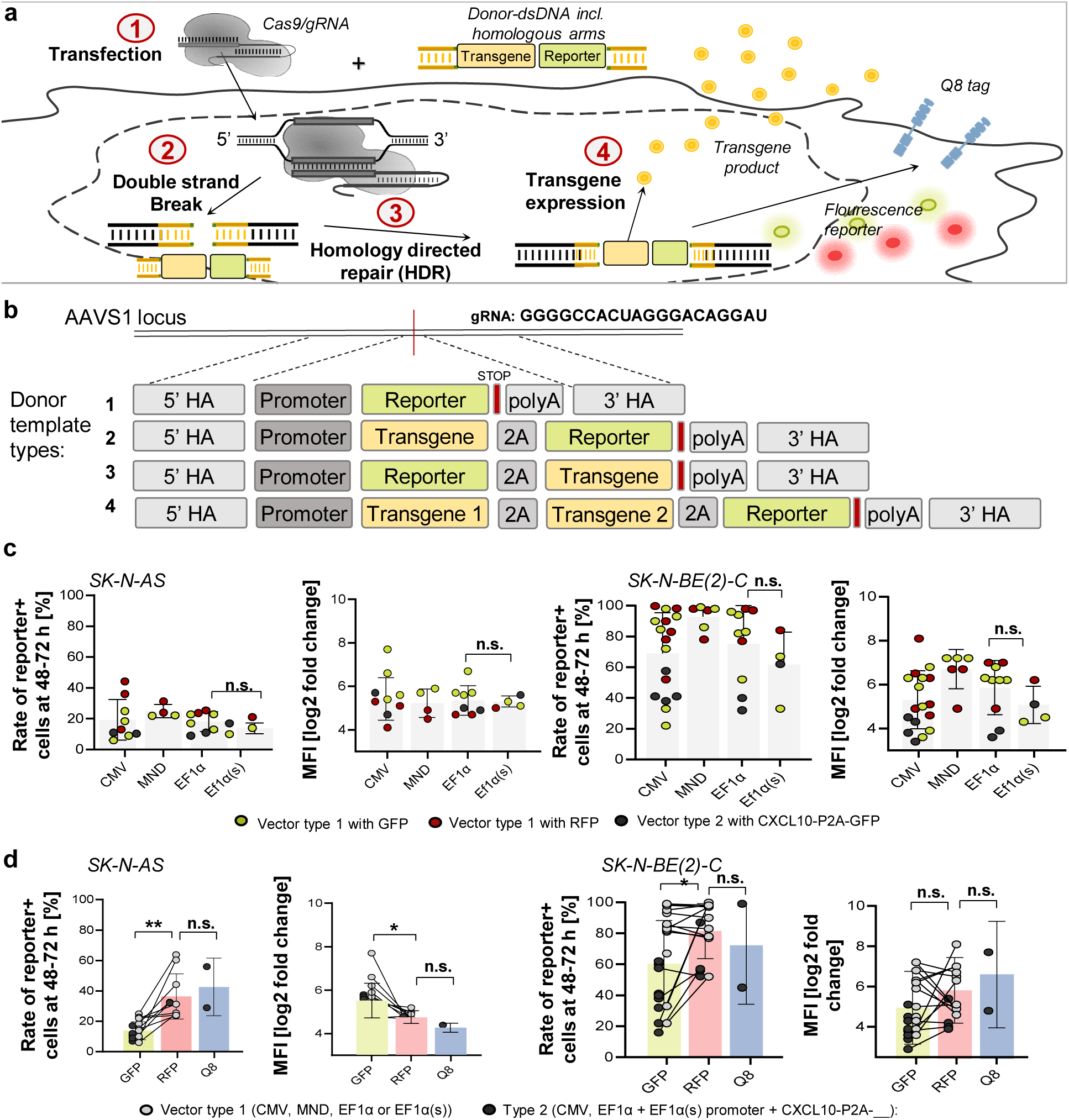
Modular CRISPR/Cas9-mediated HDR knock-in system enables virus-free transgene delivery and vector expression testing reveals cell line-dependent expression patterns in neuroblastoma cells. (a) Schematic of the CRISPR/Cas9 HDR workflow targeting the AAVS1 safe harbor locus. (1) Transfection of cells with Cas9, gRNA (RNP or mRNA/gRNA) and donor DNA. (2) Induction of double-strand breaks by Cas9. (3) Repair via HDR leading to precise integration. (4) Stable transgene expression detectable via reporter assays. (b) Modular donor designs featuring 400 bp homology arms, diverse promoters, transgenes, P2A linkers, and polyadenylation signals. Four donor template types (1–4) were tested. (c) Promoter-driven transgene expression levels in SK-N-AS and SK-N-BE(2)-C cells, showing percentages of positive cells and mean fluorescence intensity (MFI). (d) Comparison of monocistronic and bicistronic constructs encoding RFP, GFP, or Q8 reporters, illustrating differences in reporter-positive rates and MFI depending on vector design and cell line. *Data presentation: Means ± SD. Statistical analysis: (c) Mixed-effects model; (d) matched two-way ANOVA; p values: *<0.05, **<0.01; n.s., not significant*.

### CRISPR Knock-In Efficiency Depends on Construct composition and Electroporation Platform While Digital PCR Confirms Precise Genomic Integration

We systematically investigated CRISPR/Cas9-mediated knock-in performance across diverse vector architectures and delivery platforms in neuroblastoma cells. Three representative constructs of varying size and promoter context were designed: CMV-CXCL10-E2A-GFP (2010 bp insert), CMV-CXCL10-P2A-RFP (1759 bp), and EF1α(s)-CXCL10-P2A-Q8 (1065 bp), enabling assessment of size- and promoter-dependent knock-in efficiencies (Figure 2a). Flow cytometry analysis 28 days post-electroporation demonstrated an inverse relationship between construct size and stable integration rates in both neuroblastoma cell lines. EF1α(s)-CXCL10-P2A-Q8 achieved the highest integration rates (∼50% in SK-N-AS, ∼40% in SK-N-BE(2)-C), significantly exceeding the larger CMV-CXCL10-P2A-RFP (∼10%) and CMV-CXCL10-E2A-GFP (<10%) constructs (p < 0.05) (Figure 2b). Incorporation of truncated Cas9 targeting sequences (tCTS)^17^ in EF1α(s)-CXCL10-P2A-Q8 did not further enhance knock-in efficiency (Figure 2b), suggesting that construct size is the primary determinant. Digital PCR quantified integration events relative to the reference gene AFF3, distinguishing total template copy number versus precise on-target integration (Figure 2c). In line with flow cytometry data, both cell lines showed high total transgene copy number (detected by “in/in” primers) after delivery. “out/in” junction-spanning assays (dPCR and standard PCR with Sanger sequencing) confirmed site-specific knock-in on a qualitative and quantitative level. (Figure 2c, Supplementary Figure S4 and S5). Comparably lower “out/in” dPCR rates likely result from either reduced assay sensitivity due to a large amplicon size as indicated by a higher proportion of “rain” in digital PCR raw data ^40^ and/or detection of free-floating HDR templates contributing to positive signals in flow cytometry (i.e. episomal expression) or “in/in” assays. DNA dosage experiments indicated that knock-in efficiency improvements are construct-specific independently of size. For CMV_RFP-bGH (1604 bp), increasing DNA input from 1 μg to 3 μg per 10⁶ cells boosted integration from ∼30% to ∼55% (p < 0.001). However, CMV_CXCL10-RFP-sPA (1759bp) remained unaffected by higher DNA concentrations, maintaining ∼15% integration efficiency across conditions (Figure 2d). In line with previous reports, this construct-dependent dose response may reflect size- and sequence-dependent differences in donor DNA processing, where larger or more complex templates become less responsive to increased donor input due to limitations in nuclear uptake or donor stability^41,42^. Evaluation of 14 constructs revealed that the optimized construct including EF1α(s) promoter, Q8 reporter and sPA sequence yielded the strongest functional knock-in-associated flow cytometry readout across multiple designs, achieving up to 60% reporter-positive cells in SK-N-AS after 28d and substantially outperforming other constructs. The superior performance of the compact EF1α(s)-Q8-sPA cassette likely reflects a combination of reduced donor size, favorable donor architecture, and the use of regulatory and reporter elements that function efficiently in neuroblastoma cells. Smaller templates may facilitate intracellular trafficking and HDR processing, while optimized cassette composition could improve transgene expression stability and detection sensitivity. (Figure 2e). Platform benchmarking indicated minor differences between electroporation systems (Lonza 4D-Nucleofactor and Neon transfection system), with full details provided in Supplementary Note 2. Collectively, these data demonstrate that CRISPR knock-in success relies on construct size, promoter selection, and delivery platform, and that the custom designed vector including EF1α(s) promoter, Q8 reporter and sPA consistently yields superior integration efficiency across diverse experimental conditions.

**Figure 2.**
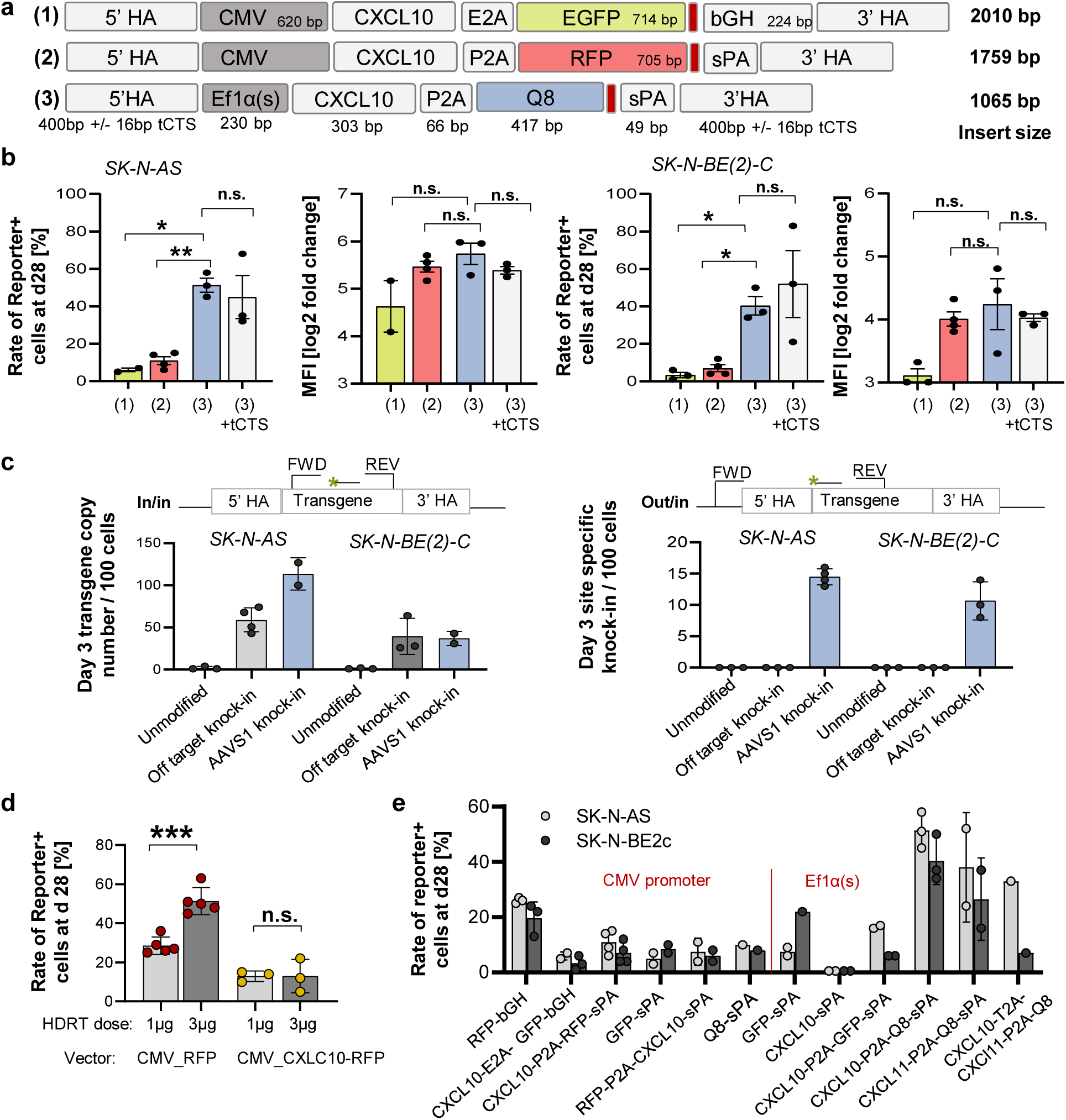
Influence of transgene size, promoter selection, and delivery platform on CRISPR/Cas9-mediated knock-in efficiency in neuroblastoma cells. (a) Schematic of three constructs differing in size and promoter context: TV2 (2010 bp), TV17 (1759 bp), and TV31 (1065 bp). All contain 5’ and 3’ homology arms for targeted integration. (b) Integration efficiency of different constructs in SK-N-AS and SK-N-BE(2)-C, measured by flow cytometry at 28 days. (c) Digital PCR quantification of total (“in/in”) versus precise on-target (“out/in”) integration. (d) DNA concentration effects on knock-in rates for TV1 and TV17. (e) Integration efficiencies across 14 different constructs highlighting superior performance of EF1α(s) promoter constructs. *Data presentation: Means ± SD. Statistical analysis: (b, d) one-way ANOVA with Dunnett’s multiple comparisons test. p-values: *<0.05, **<0.01, ***<0.001; n.s.: not significant*.

### LNP Formulation and Optimization Enables Efficient Co-Delivery of DNA and mRNA for CRISPR-Mediated Editing Despite Cell-Line Dependent Constraints

We systematically investigated LNP formulations for co-delivery of CRISPR reagents, comparing commercial PNI-LNPs (GenVoy-ILM T Cell Kit for mRNA) and self-developed ALC-LNP mixes in neuroblastoma cells (Figure 3a, b). Dynamic light scattering (DLS) analysis revealed significant differences between PNI-LNPs and ALC-LNPs in particle size, polydispersity index (PDI), and product purity. PNI-LNPs were larger (∼160 nm vs. ∼112 nm) and more heterogeneous (PDI 0.24 vs. 0.17, p < 0.0001) than ALC-LNPs, indicating better uniformity of the ALC formulations. Peak 1 area, reflecting purity, was also lower for PNI-LNPs (97.3%, p = 0.0003) compared to ∼100% for ALC-LNPs (Figure 3c, Supplementary Figure S7a). Although mRNA encapsulation efficiency was higher in PNI-LNPs (94.51%) than ALC-LNPs (88.41%; p < 0.05), both were sufficient for CRISPR applications (Figure 3c). Adjusting ALC-LNP molarity affected particle size, with lower molarity leading to increased diameters across lipid mixes A, B, and C (Figure 3d). Cell viability and proliferation were strongly dependent on LNP type and cell line. SK-N-AS cells tolerated ALC-LNPs well, maintaining growth similar to ApoE only controls (Figure 3e). In contrast, PNI-LNPs induced significant dose-dependent cytotoxicity in SK-N-AS cells, markedly reducing growth rates. (Figure 3e, f, Supplementary Figure S8a). Live-cell imaging confirmed early morphological changes in PNI-LNP-treated cells, including detachment and shrinkage by 20 hours (Supplementary Figure S8b). SK-N-BE(2)-C cells also exhibited impaired proliferation with PNI-LNPs, though the effect was less pronounced. In comparison, only weak tendencies towards reduced growth were observed for ALC-LNPs or Lipofectamine in either line (Figure 3e, f, Supplementary Figure S8b). The observed cytotoxicity was especially severe when cells were seeded at low confluence (<50%), where PNI-LNP exposure not only impaired proliferation but caused a net loss of confluence. This heightened sensitivity at low density suggests that cell-cell and cell-matrix contacts normally buffer LNP-induced membrane stress. In sparse cultures, reduced adhesion and greater exposed membrane surface may permit excessive lipid-membrane interaction, leading to pronounced cytotoxicity via physicochemical or charge-based effects. (Figure 3e, f).

**Figure 3.**
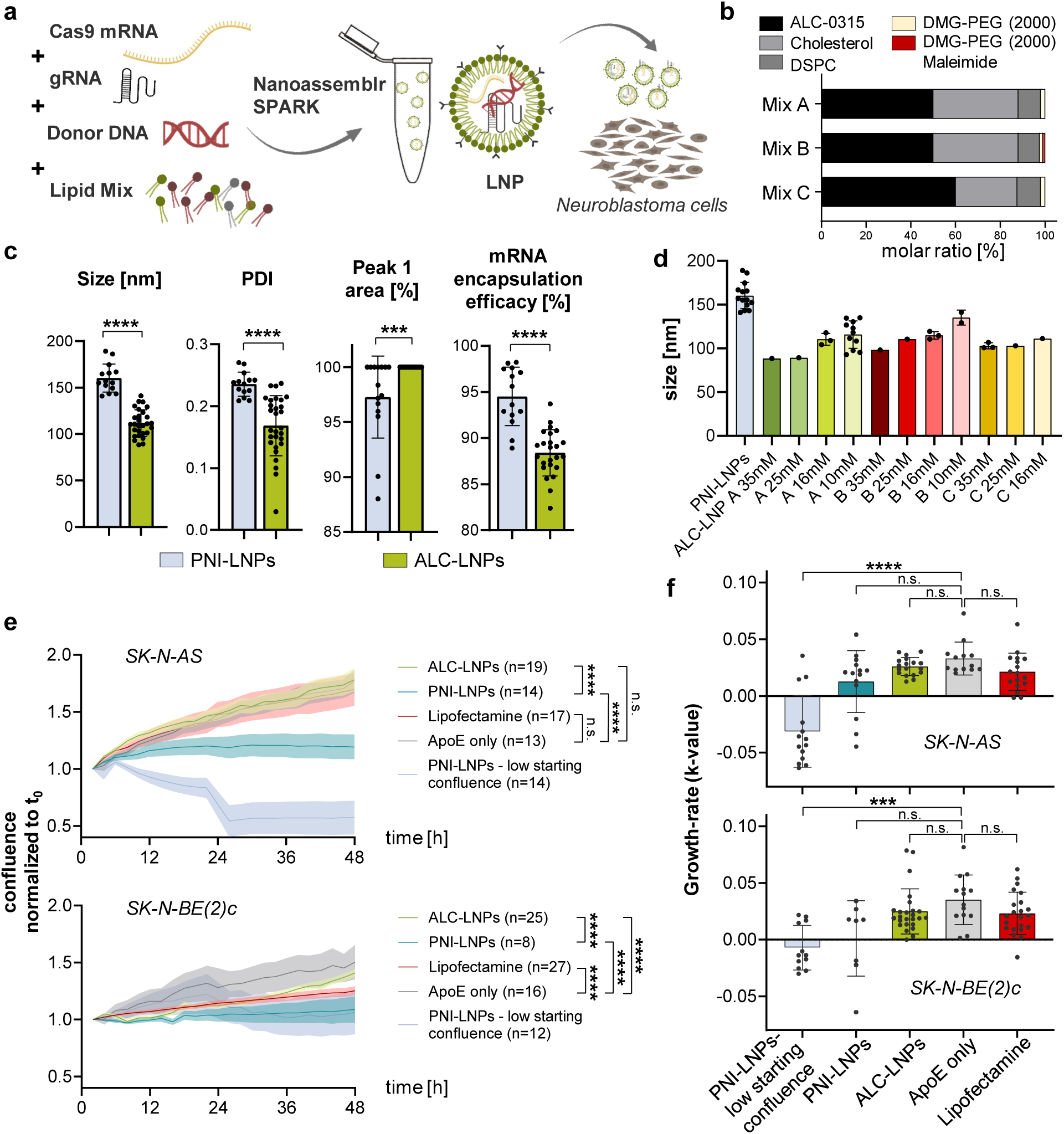
Characterization of LNP formulations and their impact on neuroblastoma cell viability. (a) Workflow for LNP preparation and application. Nucleic acid components (Cas9 mRNA, Donor DNA & gRNA) are combined with a lipid mix to formulate TME-cGT-LNPs using the Nanoassemblr SPARK, and are applied to cultured neuroblastoma cells. (b) ALC lipid mix compositions. (c) Dynamic light scattering (DLS) parameters (particle size, polydispersity index (PDI) and Peak 1 area for product purity) and mRNA encapsulation efficacy measured via ribo green assay comparing PNI-LNPs and ALC-LNPs. (d) Effect of ALC molarity on particle size. (e) Growth curves following transfection, normalized to t_0_. Initial confluence was 50–70% across conditions, except for the “PNI-LNPs – low starting confluence” group (10–50%), included to assess confluence-dependent sensitivity to LNP treatment. (f) Growth rate analysis. Data presentation: *Means ± SD. ALC-LNP data in panels c, e and f represent pooled analyses across lipid mixes A-C. Statistical analysis: (c) Mann-Whitney test, (e) Two-Way ANOVA, (f) One-Way ANOVA with Dunn’s multiple comparisons test. p-value: *<0.05, **<0.01, ***<0.001, ****<0.0001*

IncuCyte imaging for transfection efficiency analysis following co-delivery of GFP mRNA and RFP HDR templates showed that ALC-LNPs achieved the highest nucleic acid delivery in SK-N-AS cells, reaching mean RFP DNA expression of 49.7% and EGFP mRNA levels of 96.1% (Figure 4a, b). PNI-LNPs yielded intermediate results (25.5% RFP DNA, 81.6% EGFP mRNA), while Lipofectamine had the lowest efficiency (9.6% RFP DNA, 23.8% EGFP mRNA). In SK-N-BE(2)-C cells, Lipofectamine unexpectedly achieved the highest RFP DNA expression (21.09%), though this was from a single control replicate, while ALC-LNPs achieved lower RFP DNA expression (12.8%) but high EGFP mRNA delivery (94.2%) (Figure 4a, b, Supplementary Figure S9a). In this cell line, PNI-LNPs yielded the lowest RFP DNA expression (2.5%). Within ALC-LNP conditions, EGFP mRNA remained high across molarities, but RFP DNA expression was significantly higher at lower molarities, with lipid mix B at 16 mM yielding the strongest DNA signal in SK-N-AS (77.7%) and lipid mix A at 16 mM in SK-N-BE(2)-C (34.1%) (Figure 4b). Temporal analysis revealed EGFP mRNA expression appeared earlier than RFP DNA across conditions (Figure 4c). For PNI-LNPs, mRNA plateaued by 48 hours, while RFP DNA in SK-N-AS rose gradually over 96 hours. SK-N-BE(2)-C showed minimal RFP DNA expression in the first 24 hours with PNI-LNPs. In ALC-LNP-treated cells, EGFP mRNA peaked then declined, while RFP DNA continued increasing without plateau (Figure 4c). Peak RFP DNA expression times were later for ALC-LNPs (125.9 h in SK-N-AS, 131.8 h in SK-N-BE(2)-C) than for PNI-LNPs and Lipofectamine, with EGFP mRNA peaks also occurring later for ALC-LNPs (Figure 4d). The earlier peak and faster decline of EGFP mRNA compared to RFP DNA reflect fundamental differences in intracellular trafficking: mRNA typically escapes the endosome more efficiently and is translated immediately in the cytosol, whereas plasmid DNA requires nuclear entry and undergoes slower unpacking. These distinct processing routes explain the delayed and prolonged DNA expression kinetics and the cell line–dependent differences observed across LNP types. Our analysis also revealed a positive correlation between LNP size and DNA expression. Larger ALC-LNPs were associated with higher DNA expression in SK-N-AS (p = 0.0179), suggesting that increased particle size may influence DNA delivery efficiency—potentially through greater nucleic acid loading, altered endosomal trafficking, or improved intracellular stability. This trend was weaker in SK-N-BE(2)-C and should therefore be interpreted cautiously, as the relationship may be context-dependent rather than directly causal. (Figure 4e). ApoE isoform testing indicated modest differences in LNP performance between the two neuroblastoma lines (Supplementary Figure S9b, Supplementary Note 3).

**Figure 4.**
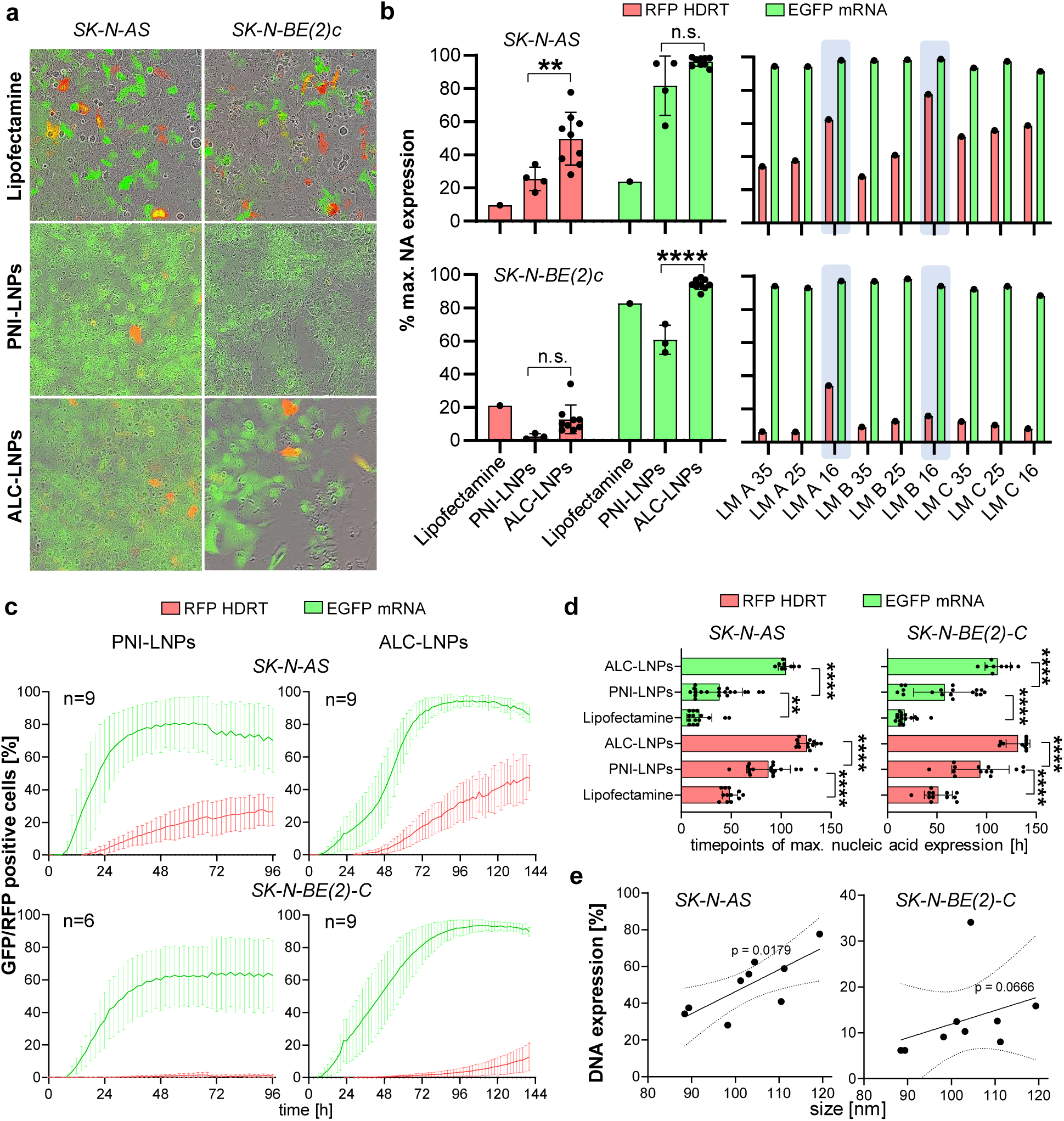
Transfection efficiency of LNPs and dynamics of nucleic acid expression. (a) Representative IncuCyte imaging of SK-N-AS and SK-N-BE(2)-C at the timepoint of max. nucleic acid expression after transfection of RFP DNA and GFP mRNA using Lipofectamine, PNI-LNPs and ALC-LNPs. (b) Max. (co-) expression of RFP DNA and EGFP mRNA comparing different transfection methods and different molarities of ALC-LNPs in SK-N-AS and SK-N-BE(2)-C as measured via IncuCyte live-cell imaging. (c) RFP DNA and EGFP mRNA expression after background substraction over 96 h measured via IncuCyte live-cell imaging. The ALC-LNP group includes both lipid mixes A and B at 16mM. (d) Time-to-peak analysis (data from (c)). (e) Correlation between particle size and DNA expression (data from (c)). *Data presentation: Means ± SD. Statistical analysis: (b, d) One-Way-ANOVA with Dunn’s; p values: **<0.01,****<0.0001; n.s.: not significant*.

Functional CRISPR editing assays using LNP delivery showed robust indel formation but lower knock-in (KI) efficiencies compared to electroporation for CMV_RFP_sPA and EF1α(s)-CXCL10-P2A-Q8-sPA repair templates. PNI-LNPs induced indels in 42.5% (SK-N-AS) and 50.5% (SK-N-BE(2)- C) initially, increasing to 88.75% and 85.5% after protocol optimization (Figure 5a, b). ALC-LNPs achieved lower indel rates of 12.5–41.0% in SK-N-AS and 25.3–68.5% in SK-N-BE(2)-C, depending on molarity and lipid mix. Despite high cutting efficiency, KI rates remained low, around ∼1.0% by dPCR out/in assays for both PNI-LNPs and ALC-LNPs (Figure 5b). The highest mean KI rate by dPCR was ∼1.4% in SK-N-BE(2)-C with ALC-LNPs lipid mix A at 10 mM, while in SK-N-AS, the maximum was ∼1.2% with PNI-LNPs. As seen with electroporation-based knock-ins, these molecular KI rates were markedly lower than flow cytometry measurements, where reporter-positive populations reached up to 18% in SK-N-BE(2)-C and 14% in SK-N-AS for PNI-LNPs, suggesting flow cytometry may detect episomal expression, and acknowledging the lower sensitivity of dPCR due to large amplicon sizes (Figure 5b). Individual replicates occasionally showed outliers with higher-than-average KI percentages, including dPCR values up to ∼5%, indicating variability due to stochastic differences in transfection efficiency, cell viability, or chromatin context (Figure 5b). Importantly, single-stranded DNA templates did not significantly improve KI rates over double-stranded DNA in either cell line, indicating template format alone was not the primary determinant of HDR success (Figure 5b). Dose comparisons showed that increasing nucleic acid from 5 μg to 17 μg per 10⁶ cells in SK-N-AS modestly improved knock-in rates, while in SK-N-BE(2)-C, higher doses gave inconsistent results and increased variability (Figure 5c). Overall, these data demonstrate that LNPs enabled efficient co-delivery of CRISPR DNA and mRNA and supported generation of genetically modified solid tumor models. While knock-in efficiencies in our experimental setting remained lower than those observed with electroporation, the successful establishment of edited cell populations highlights the potential of LNP-mediated HDR editing as a flexible non-viral platform for further development.

**Figure 5.**
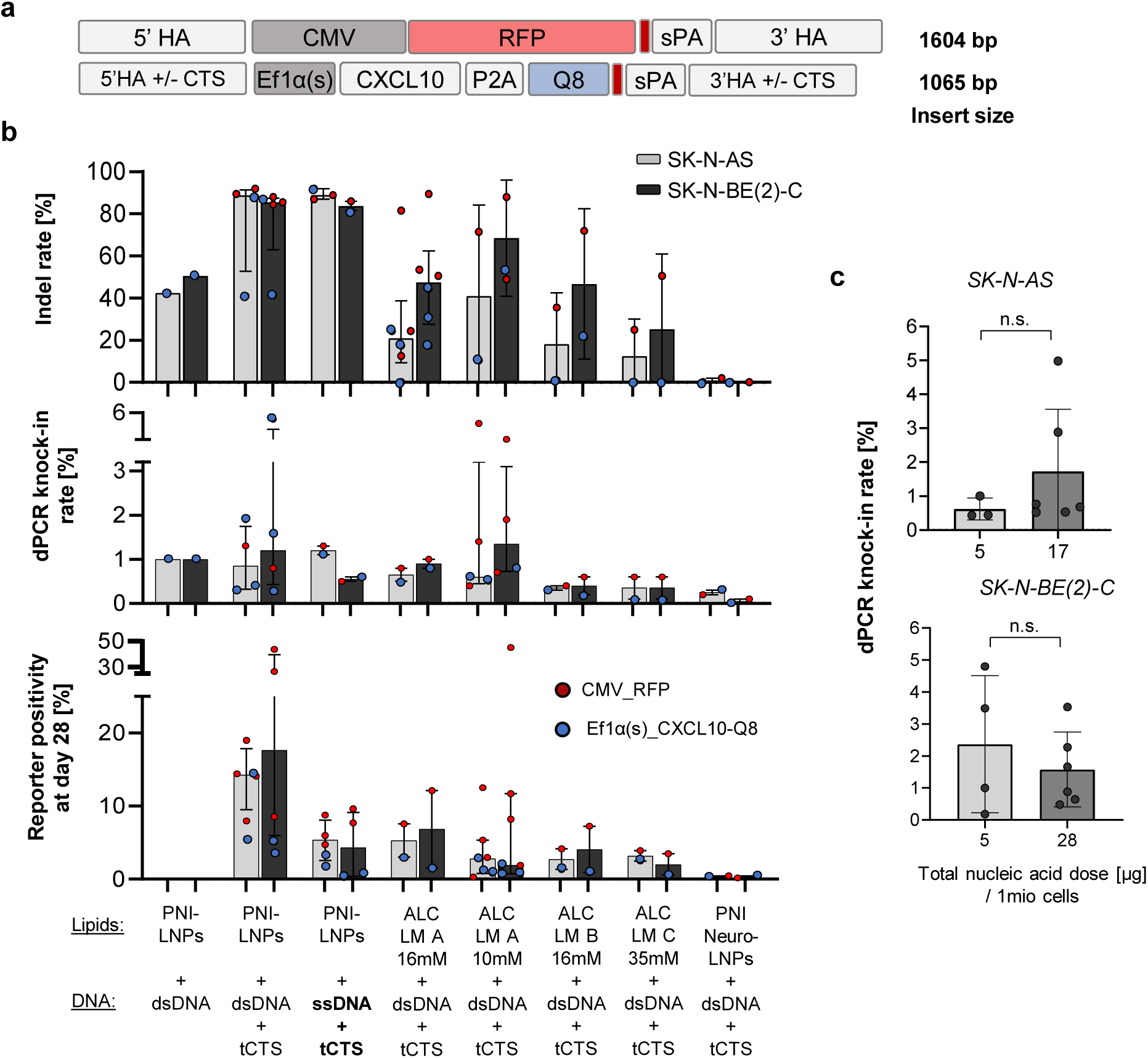
CRISPR editing efficiencies following LNP-mediated delivery. (a) HDRT designs used for LNP delivery experiments. (b) Indel rate detected by Synthego ICE analysis, knock-in rate detected using dPCR after 48h and flow cytometry data 28d after treatment. (c) Comparison of knock-in rate as measured via dPCR for different total nucleic acids dosages (mRNA+gRNA+HDRT) normalized to one million target cells from SK-N-AS or SK-N-BE(2)-C neuroblastoma cell lines. *Data presentation: Means ± SD; Statistical analysis: (c) Mann-Whitney test. P-value: n.s., not significant*.

### CRISPR Knock-In Enables Stable Chemokine Expression and Disease Modeling Using Genetically Modified Neuroblastoma Cell Lines

Building on systematic optimization of CRISPR/Cas9 knock-in efficiency, we established stable neuroblastoma cell lines harboring chemokine-reporter constructs for functional analysis and disease modeling. Polyclonal populations nucleofected with CMV_CXCL10-P2A-RFP constructs and Cas9-AAVS1-gRNA RNPs showed stable RFP positivity between 10-25% over six weeks (Figure 6a). Fluorescence-activated cell sorting (FACS) at 28 days post-transfection significantly enriched stable integrants, raising RFP-positive cells to >90% in SK-N-AS and >80% in SK-N-BE(2)-C populations (Figure 6a). Single-cell clones derived from sorted populations demonstrated high long-term stability. Functional analysis confirmed robust chemokine production from both sorted and clonal populations. ELISA measurements showed that polyclonal and clonal SK-N-AS-derived lines secreted significantly higher CXCL10 and CXCL11 levels than SK-N-BE(2)-C lines, paralleling the superior integration efficiencies in SK-N-AS cells (Figure 6b). EF1α(s)-driven constructs consistently yielded higher chemokine levels than CMV-driven constructs, reinforcing the promoter-dependent advantages established earlier (Figure 2e). Dual-expression constructs encoding EF1α(s)_CXCL10-T2A-CXCL11-P2A-Q8_sPA demonstrated simultaneous secretion of both chemokines, confirming the versatility of our modular knock-in approach for multi-gene applications (Figure 6b). We further assessed the functionality of modified cell lines in a 3D bioprinting system designed to replicate physiologically relevant tumor architecture. Modified neuroblastoma cells were successfully embedded into standardized bioprinted constructs and maintained similar viability over 96 hours regardless of genotype, as confirmed by histological analysis showing preserved morphology and even spatial distribution within the gel matrix (Figure 6c, d). CXCL10 secretion in 3D culture was consistent with levels observed in conventional 2D systems, underscoring robust transgene expression under altered physical constraints (Figure 6e). No significant differences in viability or proliferation were detected between wild-type and genetically modified cells within 3D constructs (Supplementary Figure S10). In vivo validation using a NOG xenograft model provided evidence of functional chemokine expression. Tumor growth kinetics for SK-N-AS and SK-N-BE(2)-C cells carrying integrated EF1α(s)-CXCL10-P2A-Q8 constructs were comparable to wild-type controls, indicating the genetic modifications did not impair tumorigenic capacity (Figure 6f, g, Supplementary Figure S11a). Importantly, serum CXCL10 levels were persistently elevated in mice with genetically modified tumors but remained undetectable in wild-type controls throughout the 21-day observation period, confirming sustained transgene expression under physiological conditions (Figure 6h). Flow cytometry further verified stable Q8 epitope expression post-knock-in, with modified cells showing ∼95% Q8 positivity both in vitro before transplantation and after tumor outgrowth (Supplementary Figure S11b). Morphological characterization using hematoxylin and eosin (H&E) staining revealed no discernible differences between wild-type and transgenic tumor cell lines following tumor outgrowth in vivo, indicating that CRISPR modifications did not alter basic tumor histoarchitecture (Supplementary Figure S11c). Collectively, these results demonstrate that CRISPR/Cas9-mediated knock-in enables stable transgene expression suitable for advanced in vitro and in vivo modeling applications, without detrimental effects on cellular fitness or tumorigenic potential.

**Figure 6.**
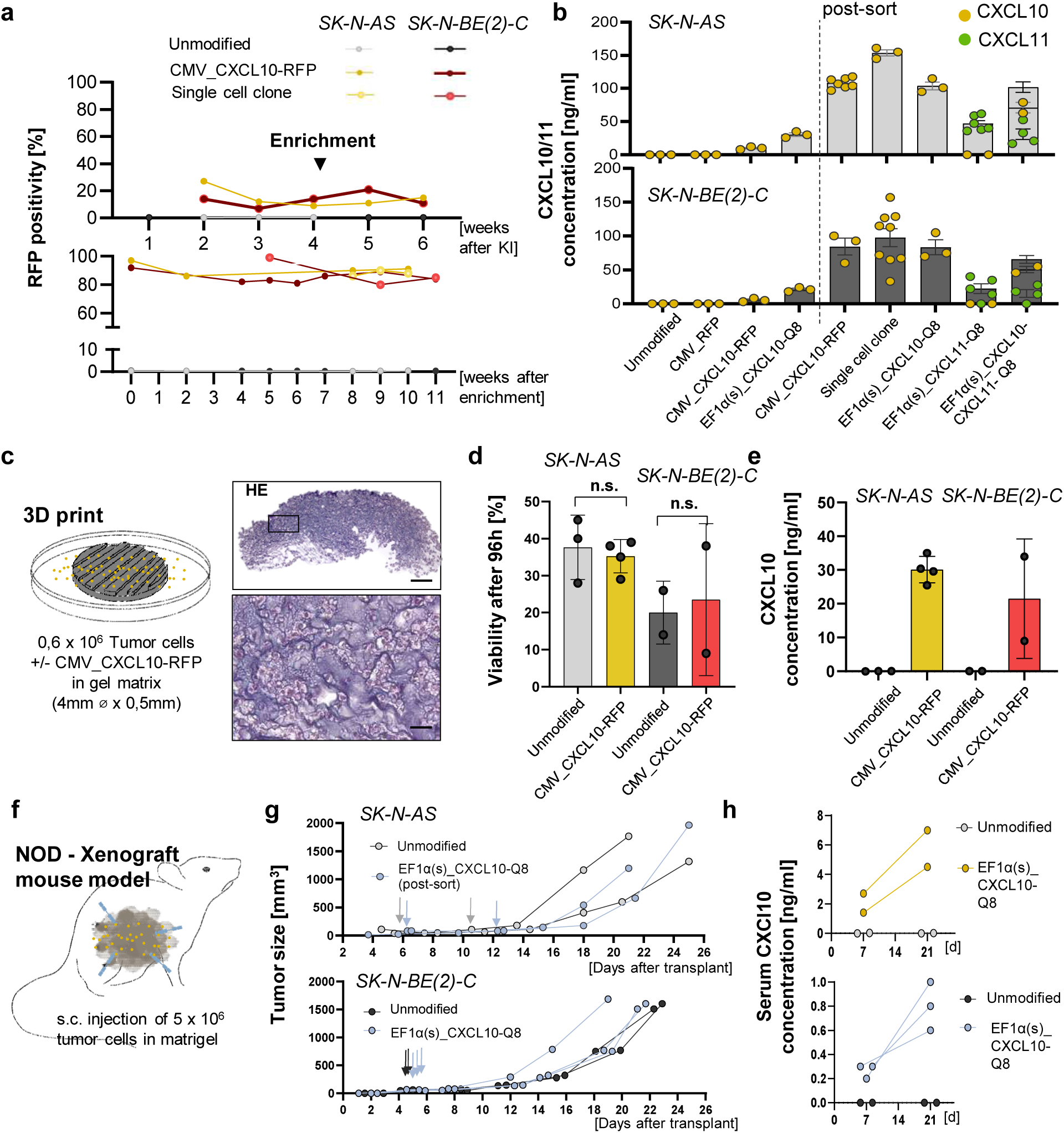
Functional validation and disease modeling using CRISPR-edited neuroblastoma cell lines. (a) Time-course analysis of RFP-positive cells (bulk or single cell) in SK-N-AS and SK-N-BE(2)-C populations pre- and post-FACS enrichment. (b) ELISA quantification of CXCL10 and CXCL11 secretion across modified cell lines before and after enrichment for transgene positive cells. 1×10^6^ tumor cells were cultivated for 72h with 1ml growth media + 10% FCS. (c) Schematic and histological analysis of 3D bioprinted constructs embedding CXCL10-expressing cells. (d) Cell viability comparison in bioprinted constructs between modified and unmodified tumor cells. (e) CXCL10 secretion levels inside of the 3D structures, measured after dissociation. (f) Schematic of NOD-SCID xenograft experiments. (g) Tumor growth kinetics in vivo. Arrows mark first time point of size >50mm^3^. (h) Serum CXCL10 levels in xenograft models. *Data presentation: Means ± SEM. Statistical analyses: (d) One-Way-ANOVA with Dunn’s multiple comparisons test; p values: n.s.: not significant*.

## Discussion

This study evaluated virus-free strategies for generating genetically modified neuroblastoma cell lines using CRISPR/Cas9-mediated homology-directed repair (HDR) as preclinical models for translational cancer research. We developed and used electroporation- and LNP-based delivery systems and investigated how construct design parameters, including size and promoter architecture, influence knock-in efficiency and stability. A notable determinant of knock-in efficiency in our study was donor construct size and design. Smaller constructs incorporating elements such as the custom designed EF1α(s) promoter and Q8 reporter consistently outperformed larger cassettes, which showed markedly reduced integration rates. This inverse relationship between donor size and HDR success has been previously documented and reflects both biophysical and cellular constraints on repair mechanisms^11,16^. In our experiments, certain constructs also demonstrated differential responsiveness to increased donor DNA amounts, suggesting that optimal donor dosage must be determined empirically for each specific design^8,15^.

Consistent with previous reports, our findings show that electroporation of Cas9 ribonucleoprotein (RNP) complexes with donor DNA achieves higher HDR-mediated knock-in efficiencies than LNP delivery, particularly for constructs below ∼2 kb in size^11,43^. While the Lonza 4D Nucleofector system provided the highest integration rates, it also resulted in somewhat reduced cell viability compared to the Neon platform. This trade-off between efficacy and cytotoxicity aligns with prior observations that the physical parameters of electroporation can induce varying degrees of cell stress^14^. LNP-based delivery showed lower HDR efficiency than electroporation, despite achieving high levels of mRNA delivery and indel formation. Even under optimized conditions, HDR knock-in rates remained generally below ∼1–2% by digital PCR. These results are consistent with other studies indicating challenges in achieving efficient nuclear localization of HDR templates with LNP-based delivery necessary for efficient knock-in^44,45^. Nevertheless, our data confirm that even modest HDR efficiencies from LNP delivery can be practically useful when coupled with enrichment methods such as fluorescence-activated cell sorting (FACS). Post-sorting populations in our experiments reached transgene-positive fractions exceeding 80%, demonstrating the feasibility of non-viral CRISPR-based genome editing for generating stable cell lines despite low initial knock-in rates. In contrast to electroporation, LNP-based delivery may additionally offer future compatibility with in vivo genome engineering approaches, potentially increasing the translational relevance of tumor models generated using adaptable non-viral HDR workflows. Further on, cell line-specific differences significantly influence nucleic acid expression and HDR rates. These differences arise from intrinsic variations in genetic composition, metabolic capacity, and signaling networks, resulting in divergent responses even under identical experimental conditions. Consequently, protocols optimized for one cellular model may not apply to another, making context-dependent optimization crucial for reproducible and valid outcomes across various applications. Another consideration in CRISPR knock-in workflows is the time required to establish pure transgenic populations. Traditional approaches necessitate several weeks of culture to allow dilution of episomal expression and ensure stable integration before enrichment. Strategies such as in-frame knock-ins, where transgenes are fused directly to endogenous coding regions, could help identify successfully modified cells earlier and reduce overall timelines. This approach also maintains endogenous regulatory context, potentially preserving physiological gene expression levels and reducing artifacts associated with exogenous promoters^1,46^.

Looking forward, emerging genome editing technologies may help address the current limitations of HDR-based knock-in. Prime editing offers the ability to introduce precise sequence changes without double-strand breaks but remains less efficient for large insertions^22^. CRISPR-associated transposase systems show potential for programmable integration of larger DNA fragments but are still under development for practical applications^24^. Until these approaches mature, further refinement of HDR protocols - including construct engineering, delivery optimization, and post-editing enrichment - remains critical for advancing non-viral genome engineering. Taken together, our findings highlight both the potential and the constraints of virus-free CRISPR knock-in approaches for generating stable, genetically modified tumor cell lines. In our work electroporation is the most effective method for achieving high HDR efficiency, particularly for small to moderate-sized constructs. However, advances in LNP technology and peptide-based systems and vector design are expanding the feasibility of non-viral delivery systems^25,26,47–49^. Our results further indicate that custom designed non-proprietary ALC-based formulations can support CRISPR cargo delivery and HDR-associated editing in tumor cells, offering a flexible framework for future optimization of non-viral delivery systems. These developments are particularly relevant for applications such as immunotherapy research, where virus-free non-toxic engineering strategies may reduce confounding factors, safety risks and simplify regulatory pathways^20,21^. Further work should focus on improving knock-in efficiencies for larger constructs, minimizing cellular toxicity, for example through developing more efficient LNP platforms or testing other delivery systems like peptide-based delivery approaches^48,49^. Additionally, evaluating the long-term genomic stability and functional consequences of knock-ins in different cellular contexts remains an important area for future investigation.

## Author contributions

SA, MP, JM, EO, LR, CV, CL, MS and ML performed the experiments. SA, MP and ML analyzed the data.

SA, MP, CV, JK, VG, AN, MS, ND, KA, DLW, RK, AK and ML developed the methodology.

SA, MP and ML wrote the manuscript with input from all authors.

RK, AK and ML, supervised the study.

AK and ML acquired funding for the study.

ML conceived and designed the study.

## Data sharing statement

All data collected and analyzed in this study as well as detailed test statistics can be received upon request. This excludes data that falls under confidentiality restrictions because it would allude to unpublished data of other projects. Step-by-step protocols generated in this study, including interactive setup sheets for CRISPR knock-in workflows, will be made available in protocols.io upon publication.

## Conflict of interest disclosure

ML has received reagents and services related to CRISPR-Cas gene editing and digital PCR from Qiagen. DLW is named as an inventor on patent applications related to genome editing and cell therapies. DLW is a co-founder and holds equity in TCBalance Biopharmaceuticals GmbH. DLW’s laboratory at Charité has received reagents and services related to CRISPR-Cas gene editing from Integrated DNA Technologies and GenScript Inc.

None of the aforementioned companies were involved in the design, execution, or interpretation of the study. All other authors declare no competing interests.

## Declaration of Generative AI and AI-assisted technologies in the writing process

During the preparation of this work the authors used ChatGPT (https://chat.openai.com/) by OpenAI (2026) in order to restructure and refine sentences and paragraphs for clarity and flow. After using this tool/service, the authors reviewed and edited the content as needed and take full responsibility for the content of the publication.

## Acknowledgments

The authors would like to thank Silke Schwiebert and Anika Winkler for their technical support in conducting the experiments, and Angelika Eggert and Nikolaus Rajewsky for their support as mentors within the framework of the BIH Charité Clinician Scientist Program.

MP gratefully acknowledges the support of a doctoral scholarship provided by the Jürgen Manchot Stiftung. SA gratefully acknowledges funding from the Berlin Institute of Health (BIH) through a doctoral fellowship. This work was supported by the Berliner Krebsgesellschaft (grant number LAFF202008 to M.L.) and KINDerLEBEN e.V. Berlin (to M.L.). Additional support was provided by Charité – Universitätsmedizin Berlin and the Berlin Institute of Health (BIH) at Charité – Universitätsmedizin Berlin (AdHoc Booster Grant to D.W. and M.L.) and the DZG Innovation Fund (to A.K.).

M.L. participates in the BIH Charité Clinician Scientist Program, funded by Charité – Universitätsmedizin Berlin and the BIH. A.K. participates in the BIH Charité Advanced Clinician Scientist Pilot Program, also funded by Charité – Universitätsmedizin Berlin and the BIH.

## Notes

### Competing Interest Statement

The authors have declared no competing interest.

## References

1. Wang, H., La Russa, M. & Qi, L. S. CRISPR/Cas9 in Genome Editing and beyond. Annu. Rev. Biochem. 85, 227–264 (2016).

2. Kath, J. et al. Pharmacological interventions enhance virus-free generation of TRAC-replaced CAR T cells. Mol. Ther. Methods Clin. Dev. 25, 311–330 (2022).

3. Milone, M. C. & O’Doherty, U. Clinical use of lentiviral vectors. Leukemia 32, 1529–1541 (2018).

4. Themis, M. et al. Oncogenesis following delivery of a nonprimate lentiviral gene therapy vector to fetal and neonatal mice. Molecular Therapy 12, 763–771 (2005).

5. Chandler, R. J. et al. Vector design influences hepatic genotoxicity after adeno-associated virus gene therapy. Journal of Clinical Investigation 125, 870–880 (2015).

6. Wang, Y. et al. Suicidal Autointegration of Sleeping Beauty and piggyBac Transposons in Eukaryotic Cells. PLoS Genet. 10, (2014).

7. Dunbar, C. E. et al. Gene therapy comes of age. Science (1979). 359, (2018).

8. Liao, H., Wu, J., VanDusen, N. J., Li, Y. & Zheng, Y. CRISPR-Cas9-mediated homology-directed repair for precise gene editing. Mol. Ther. Nucleic Acids 35, (2024).

9. Cong, L. et al. Multiplex genome engineering using CRISPR/Cas systems. Science (1979). 339, 819–823 (2013).

10. Mali, P. et al. RNA-guided human genome engineering via Cas9. Science (1979). 339, 823–826 (2013).

11. Richardson, C. D., Ray, G. J., DeWitt, M. A., Curie, G. L. & Corn, J. E. Enhancing homology-directed genome editing by catalytically active and inactive CRISPR-Cas9 using asymmetric donor DNA. Nat. Biotechnol. 34, 339–344 (2016).

12. Ghanta, K. S. et al. 5′ modifications improve potency and efficacy of dna donors for precision genome editing. Elife 10, (2021).

13. Jin, Y. Y. et al. Enhancing homology-directed repair efficiency with HDR-boosting modular ssDNA donor. Nat. Commun. 15, (2024).

14. Paquet, D. et al. Efficient introduction of specific homozygous and heterozygous mutations using CRISPR/Cas9. Nature 533, 125–129 (2016).

15. Carlson-Stevermer, J. et al. Assembly of CRISPR ribonucleoproteins with biotinylated oligonucleotides via an RNA aptamer for precise gene editing. Nat. Commun. 8, (2017).

16. Bae, S., Kweon, J., Kim, H. S. & Kim, J. S. Microhomology-based choice of Cas9 nuclease target sites. Nat. Methods 11, 705–706 (2014).

17. Nguyen, D. N. et al. Polymer-stabilized Cas9 nanoparticles and modified repair templates increase genome editing efficiency. Nat. Biotechnol. 38, 44–49 (2020).

18. Amiri, M., Moaveni, A. K., Majidi Zolbin, M., Shademan, B. & Nourazarian, A. Optimizing cancer treatment: the synergistic potential of CAR-T cell therapy and CRISPR/Cas9. Front. Immunol. 15, (2024).

19. Chiesa, R. et al. Universal Base-Edited CAR7 T Cells for T-Cell Acute Lymphoblastic Leukemia. New England Journal of Medicine 10.1056/NEJMOA2505478 (2025) doi:10.1056/NEJMOA2505478.

20. Song, P. et al. CRISPR/Cas-based CAR-T cells: production and application. Biomark. Res. 12, (2024).

21. Azeez, S. S. et al. Advancing CAR-based cell therapies for solid tumours: challenges, therapeutic strategies, and perspectives. Mol. Cancer 24, 191 (2025).

22. Anzalone, A. V. et al. Search-and-replace genome editing without double-strand breaks or donor DNA. Nature 576, 149–157 (2019).

23. Anzalone, A. V., Koblan, L. W. & Liu, D. R. Genome editing with CRISPR–Cas nucleases, base editors, transposases and prime editors. Nat. Biotechnol. 38, 824–844 (2020).

24. Klompe, S. E., Vo, P. L. H., Halpin-Healy, T. S. & Sternberg, S. H. Transposon-encoded CRISPR–Cas systems direct RNA-guided DNA integration. Nature 571, 219–225 (2019).

25. Chen, K. et al. Lung and liver editing by lipid nanoparticle delivery of a stable CRISPR–Cas9 ribonucleoprotein. Nat. Biotechnol. 10.1038/S41587-024-02437-3, (2024) doi:10.1038/S41587-024-02437-3,.

26. Lee, K. et al. Nanoparticle delivery of Cas9 ribonucleoprotein and donor DNA in vivo induces homology-directed DNA repair. Nat. Biomed. Eng. 1, 889–901 (2017).

27. Liu, C., Zhang, L., Liu, H. & Cheng, K. Delivery strategies of the CRISPR-Cas9 gene-editing system for therapeutic applications. Journal of Controlled Release 266, 17–26 (2017).

28. Gillmore, J. D. et al. CRISPR-Cas9 In Vivo Gene Editing for Transthyretin Amyloidosis. New England Journal of Medicine 385, 493–502 (2021).

29. Rurik, J. G. et al. CAR T cells produced in vivo to treat cardiac injury. Science (1979). 375, 91–96 (2022).

30. Kosicki, M., Tomberg, K. & Bradley, A. Repair of double-strand breaks induced by CRISPR–Cas9 leads to large deletions and complex rearrangements. Nat. Biotechnol. 36, 765–771 (2018).

31. Haapaniemi, E., Botla, S., Persson, J., Schmierer, B. & Taipale, J. CRISPR-Cas9 genome editing induces a p53-mediated DNA damage response. Nat. Med. 24, 927–930 (2018).

32. Akinc, A. et al. Targeted Delivery of RNAi Therapeutics With Endogenous and Exogenous Ligand-Based Mechanisms. Molecular Therapy 18, 1357 (2010).

33. Kath, J. et al. Integration of ζ-deficient CARs into the CD3ζ gene conveys potent cytotoxicity in T and NK cells. Blood 143, 2599–2611 (2024).

34. Nitulescu, A. M. et al. Single-stranded HDR templates with truncated Cas12a-binding sequences improve knock-in efficiencies in primary human T cells. Mol. Ther. Nucleic Acids 36, 102568 (2025).

35. Glaser, V. et al. Combining different CRISPR nucleases for simultaneous knock-in and base editing prevents translocations in multiplex-edited CAR T cells. Genome Biol. 24, (2023).

36. Rosenblum, D. et al. CRISPR-Cas9 genome editing using targeted lipid nanoparticles for cancer therapy. Sci. Adv. 6, (2020).

37. Kenjo, E. et al. Low immunogenicity of LNP allows repeated administrations of CRISPR-Cas9 mRNA into skeletal muscle in mice. Nat. Commun. 12, (2021).

38. Grunewald, L. et al. A Reproducible Bioprinted 3D Tumor Model Serves as a Preselection Tool for CAR T Cell Therapy Optimization. Front. Immunol. 12, (2021).

39. Bister, A. et al. A novel CD34-derived hinge for rapid and efficient detection and enrichment of CAR T cells. Mol. Ther. Oncolytics 23, 534–546 (2021).

40. Huggett, J. F., Cowen, S. & Foy, C. A. Considerations for digital PCR as an accurate molecular diagnostic tool. Clin. Chem. 61, 79–88 (2015).

41. Shy, B. R. et al. High-yield genome engineering in primary cells using a hybrid ssDNA repair template and small-molecule cocktails. Nat. Biotechnol. 41, 521–531 (2023).

42. Lukacs, G. L. et al. Size-dependent DNA mobility in cytoplasm and nucleus. J. Biol. Chem. 275, 1625–1629 (2000).

43. Lee, J. S., Kallehauge, T. B., Pedersen, L. E. & Kildegaard, H. F. Site-specific integration in CHO cells mediated by CRISPR/Cas9 and homology-directed DNA repair pathway. Sci. Rep. 5, (2015).

44. Wilson, R. C. & Gilbert, L. A. The Promise and Challenge of in Vivo Delivery for Genome Therapeutics. ACS Chem. Biol. 13, 376–382 (2018).

45. Mitchell, M. J. et al. Engineering precision nanoparticles for drug delivery. Nat. Rev. Drug Discov. 20, 101–124 (2021).

46. Leonetti, M. D., Sekine, S., Kamiyama, D., Weissman, J. S. & Huang, B. A scalable strategy for high-throughput GFP tagging of endogenous human proteins. Proc. Natl. Acad. Sci. U. S. A. 113, E3501–E3508 (2016).

47. Cheng, Q. et al. Selective organ targeting (SORT) nanoparticles for tissue-specific mRNA delivery and CRISPR–Cas gene editing. Nat. Nanotechnol. 15, 313–320 (2020).

48. Foss, D. V. et al. Peptide-mediated delivery of CRISPR enzymes for the efficient editing of primary human lymphocytes. Nat. Biomed. Eng. 7, 647–660 (2023).

49. Sahu, S. U. et al. Peptide-enabled ribonucleoprotein delivery for CRISPR engineering (PERC) in primary human immune cells and hematopoietic stem cells. Nat. Protoc. 10.1038/S41596-025-01154-8, (2025) doi:10.1038/S41596-025-01154-8,.

