## Supplemental file for "A Non-Viral CRISPR/Cas9 HDR Platform for Stable Engineering of Solid Tumor Models"

- Supplement -

**Content**

### List of Abbreviations

AAV - Adeno-associated virus

AAVS1 - Adeno-associated virus integration site 1

AE - Elution buffer AE

ALC - Ionizable lipid component ALC-0315

ANOVA - Analysis of variance

APC - Allophycocyanin

AS - SK-N-AS neuroblastoma cell line

ATCC - American Type Culture Collection

bGH - Bovine growth hormone polyadenylation signal

BHQ1 - Black Hole Quencher 1

CAR - Chimeric antigen receptor

cDNA - Complementary DNA

CMV - Cytomegalovirus promoter

CRISPR - Clustered regularly interspaced short palindromic repeats

CXCL10 - C-X-C motif chemokine ligand 10

CXCL11 - C-X-C motif chemokine ligand 11

dPCR - Digital polymerase chain reaction

DLS - Dynamic light scattering

dsDNA - Double-stranded DNA

E2A - Thosea asigna virus 2A peptide

EF1α - Elongation factor 1 alpha promoter

EF1α(s) - Shortened EF1α promoter

EGFP - Enhanced green fluorescent protein

ELISA - Enzyme-linked immunosorbent assay

FACS - Fluorescence-activated cell sorting

FMO - Fluorescence minus one

GFP - Green fluorescent protein

gRNA - Guide RNA

H& - Hematoxylin and eosin

HDR- Homology-directed repair

HDRT - Homology-directed repair template

ICE - Inference of CRISPR Edits

IL-15- Interleukin-15

LNP - Lipid nanoparticle

MFI - Mean fluorescence intensity

MND - Myeloproliferative sarcoma virus enhancer promoter

NSG - NOD-scid IL2Rγnull

P2A - Porcine teschovirus 2A peptide

PAM - Protospacer adjacent motif

PDI - Polydispersity index

PNI - Precision Nanosystems Inc. (GenVoy ILM)

qPCR - Quantitative polymerase chain reaction

Q8 - Custom CD34 epitope-based reporter tag

RFP - Red fluorescent protein

RNP - Ribonucleoprotein (Cas9-gRNA complex)

RPM - Revolutions per minute

SD - Standard deviation

sgRNA - Single-guide RNA

sPA - Synthetic polyadenylation signal

ssDNA - Single-stranded DNA

T2A - Thosea asigna virus 2A peptide

TME - Tumor microenvironment

UTR - Untranslated region

### Supplementary Methods

#### ssDNA Production

To produce single strand DNA (ssDNA), we used the Guide-it™ Long ssDNA Production System v2 (Takara). First, a PCR (KAPA HiFi HotStart 2x ReadyMix (Roche)) was performed, using phosphorylated primers for one strand and non-phosphorylated primers for the other. The dsDNA yield was cleaned up using AMPure XP beads (Beckman Coulter) and 70% ethanol, eluted in 50 µl nfH2O (from the kit) and quantified using Nanodrop. For ssDNA generation, 10-15 µg dsDNA were added to 5 µl strandase buffer (10x), 10µl digestion enhancer (5x) and 5 µl strandase mix A, then nfH2O from the kit was added up to a volume of 50 µl. After vortexing the reaction mix, it was incubated at 37°C for 5 min/kb, then 5 minutes at 80°C and then let cool down for 2-3 minutes on ice. In the next step, 1 µl/reaction strandase B was added. Again, the mix was incubated at 37°C for 5 min/kb, then 5 minutes at 80°C and then put on ice to cool down until clean up. Finally, a second clean up step was performed using AMPure XP beads, 70% ethanol and a magnet stand. This time, only 15 µl ddH2O was used for elution, ssDNA concentration was measured using a NanoDrop 2000 spectrophotometer (Thermo Fisher Scientific).

#### Vector Expression Testing Protocol Using Effectene Transfection

Cells were plated in 12-well plates to reach 40-80% confluence on the day of transfection. SK-N-BE(2)-C cells were seeded at 1 × 10^5^ cells per well, and SK-N-AS cells at 1.3 × 10^5^ cells per well. Confluence was confirmed visually, then 750 µL of fresh medium was added to each well. For each transfection, 40 µL EC-Buffer (QIAGEN), 2 µL Effectene Enhancer (QIAGEN), and 200 ng of plasmid DNA (~5 kb vector), HDRT, mRNA, or sgRNA were combined and incubated at room temperature for 5 minutes. Next, 2 µL Effectene Transfection Reagent was added to the mixture to form DNA-reagent complexes. Cells were washed with PBS, and the complexes were applied dropwise to the wells. All transfections were performed in triplicate. At 24 hours post-transfection, transfection complexes were removed by washing cells with PBS, and fresh medium was added. Transient expression was monitored using either live imaging (Incucyte, Sartorius) or flow cytometry. For flow cytometric analysis, transgene expression was assessed 48-72 hours after transfection.

#### Lipofectamine™ 3000 & Lipofectamine™ MessengerMax™

Neuroblastoma cells were seeded and incubated 24 hours in advance in RPMI 1640 media (Gibco™) with 10% FCS and penicillin-streptomycin (Gibco™), aiming for 70-90% confluency in a 96 well plate at the time of transfection. Before treatment we washed the cells and added 750µl fresh medium. We transfected HDRT, mRNA and sgRNA in three consecutive steps, using Lipofectamine™ 3000 for DNA and sgRNA and MessengerMAX™ (Invitrogen™) for mRNA with optimized manufacturer protocols. Reagents of each step were mixed on a separate plate and then applied dropwise onto the sample wells. After transfection, the plates were transferred to the Incucyte® (Sartorius) to monitor expression.

#### Live-Cell Imaging and Analysis

We obtained data on cell growth, toxicity and nucleic acid expression through Live-cell imaging using the Incucyte® S3 from Sartorius. For imaging, all cell lines were kept in RPMI 1640 (Gibco) or ImmunoCult TM-XF T Cell expansion medium to optimize image quality. The cells were imaged in 2-hour intervals over different periods.

#### Genomic DNA Isolation, PCR Amplification, and Purification

Genomic DNA was extracted from cultured or treated cells using the QIAamp DNA Mini Kit (QIAGEN) following the manufacturer’s instructions with slight optimizations. Typically, up to 5 × 10⁶ cells were lysed with Proteinase K and 200 μL Buffer AL at 56°C for 10 minutes. After addition of 200 μL ethanol, the lysate was loaded onto QIAamp Mini Spin Columns and centrifuged at 6,000 × g (8,000 rpm) for 1 minute to bind DNA. Bound DNA was washed sequentially with 500 μL Buffer AW1 (centrifuged at 6,000 × g) and Buffer AW2 (centrifuged at 20,000 × g). DNA was eluted with 200 μL of either Buffer AE or double-distilled water (ddH₂O) after a 1-minute incubation at room temperature, followed by a final centrifugation at 6,000 × g. DNA concentration and purity were assessed by measuring absorbance at 260 nm using a NanoDrop spectrophotometer (Thermo Fisher Scientific), ensuring yields of at least 10 ng/μL for downstream applications. Polymerase chain reactions (PCR) were carried out using KAPA polymerase (Roche) to amplify DNA fragments for homology-directed repair templates (HDRTs), target region sequencing, and qualitative knock-in analysis. Primers were custom-designed with SnapGene (GSL Biotech LLC), Primer3Plus (https://www.primer3plus.com/), and NCBI Primer-BLAST. Reactions were performed on a Bio-Rad C1000 Touch Thermal Cycler under optimized thermal cycling conditions. PCR products were purified using the QIAquick PCR Purification Kit (QIAGEN). For purification, five volumes of Buffer PB were mixed with one volume of PCR product, loaded onto the provided columns, and centrifuged at 17,900 × g (13,000 rpm) for 1 minute. Columns were washed with 750 μL Buffer PE, and DNA was eluted with 50 μL ddH₂O following a 1-minute incubation at room temperature and centrifugation. Purified PCR products were quantified on a NanoDrop spectrophotometer. Amplicon specificity and size were confirmed via electrophoresis on 1-2% agarose gels in TAE buffer stained with GelRed (Sigma-Aldrich). Sequencing of purified PCR products was performed by LGC Genomics for validation. A detailed list of all primers and oligonucleotides used is provided in **Supplementary Tables 2 - 4**.

#### gRNA Cutting Efficiency Testing

sgRNAs were designed using online CRISPR tools and synthesized by Synthego. CRISPR editing efficiency for gRNAs targeting the AAVS1 safe harbor locus was evaluated in a five-step workflow: sgRNA design, RNP complex nucleofection or LNP Cas mRNA and sgRNA cotransfection, sample preparation, target region amplification, and quantification of editing outcomes. A target-specific PCR assay was created with GeneGlobe (QIAGEN). RNP complexes were delivered into cells via the Lonza 4D Nucleofection system with or without HDR templates and Cas mRNA and sgRNA with or without HDR templates via LNPs. Treated cells were seeded into 96-well plates and cultured for 48 h at 37°C, 5% CO₂. Editing efficiency was quantified by PCR amplification of regions flanking the CRISPR target site (KAPA HiFi polymerase), followed by Sanger sequencing. Indel frequencies were analyzed using the Synthego ICE software.

#### Genome-wide Copy Number Variation (CNV) Analysis by Hi-C

Genome-wide CNV analysis of wild-type SK-N-AS and SK-N-BE2c neuroblastoma cell lines was performed using the Hi-C method to detect large-scale structural variations. Hi-C libraries were prepared following the QIAGEN EpiTect® Hi-C Kit protocol. Briefly, adherent cells were harvested by trypsinization, washed with PBS, and crosslinked with 37% formaldehyde. Crosslinking was quenched by adding 3 M Tris (pH 7.5), followed by cell lysis. Chromatin digestion was carried out using a restriction enzyme that generates 5′ overhangs, which were subsequently filled in with biotinylated nucleotides. Proximity ligation was performed under dilute conditions to favor intramolecular ligations between crosslinked DNA fragments. After reverse crosslinking and protein digestion with Proteinase K, ligated DNA was purified, fragmented via sonication (Covaris S220), and biotinylated fragments were isolated using streptavidin-coated magnetic beads. End repair, A-tailing, and adapter ligation were performed to generate sequencing-ready libraries, which were amplified by PCR and purified. Libraries were quality-checked using an Agilent 2100 Bioanalyzer and quantified by qPCR (QIAseq Library Quant Kit, QIAGEN). Paired-end sequencing was conducted on an Illumina platform (NextSeq 500/550 v2.5). For CNV analysis, sequencing data were processed to create Hi-C contact matrices. Copy number estimation was performed using the HiNT pipeline, which calculates log2 ratios of observed read coverage to expected coverage across genomic bins. Profiles were visualized as genome-wide scatter plots to identify gains and losses in chromosomal regions. CNV plots for SK-N-AS and SK-N-BE2c wild-type cells were generated, revealing characteristic patterns of chromosomal copy number alterations associated with neuroblastoma cell lines.

#### Validation of Transgene Knock-In

To confirm both efficiency and site-specificity of CRISPR/Cas-mediated knock-in, we employed three orthogonal assays: (i) flow cytometry for reporter or epitope-tag expression, (ii) junction-spanning standard PCR with Sanger sequencing to verify targeted integration, and (iii) digital PCR (dPCR) for quantification. Flow cytometric analyses were performed on a BD LSRFortessa X-20 (lasers: 405, 488, 561, 640 nm). For fluorescent reporters (e.g., eGFP, GFP, RFP), cells were harvested at defined time points post-nucleofection, washed twice with PBS, and analyzed directly using appropriate laser/emission settings (e.g., 488 nm excitation, 530 ± 30 nm emission for eGFP). For epitope detection (e.g., Q8), cells were stained in 100 μL PBS with PE-conjugated anti-CD34 antibody (clone QBEnd/10, Invitrogen; 1:10 dilution) for 30 min at 4°C in the dark, washed twice (1,300 rpm, 3 min), and resuspended in PBS for immediate analysis. Viability was assessed using fixable viability dye APC-Cy7 (1:5000, Thermo Fisher Scientific). Data were analyzed in FlowJo v10, with knock-in efficiency reported as the percentage of fluorescent or CD34-positive cells within the live population. For enrichment, CD34-positive cells were sorted on a BD FACSAria III into FBS-coated tubes, expanded in culture, and confirmed for post-sort purity by reanalysis. Standard PCR and Sanger Sequencing: To confirm site-specific integration, junction-spanning PCR assays were performed using primers designed in NCBI Primer-BLAST, Primer3Plus, and SnapGene. For each junction, one primer annealed to genomic DNA beyond the homology arm, and the other to the inserted cassette, ensuring amplification only upon correct HDR-mediated integration. All primer sequences and coordinates are listed in **Supplementary Tables 2 - 3**. Genomic DNA was extracted from cells using the QIAamp DNA Mini Kit (QIAGEN). Up to 5 × 10⁶ cells were lysed in Buffer AL with Proteinase K at 56°C for 15 min, followed by ethanol precipitation, silica column purification, and elution in 100 μL Buffer AE. DNA yield and purity were measured on a NanoDrop 2000 spectrophotometer (Thermo Fisher Scientific). PCR reactions were carried out with either Q5® Hot Start High-Fidelity 2× Master Mix (NEB) or KAPA HiFi HotStart ReadyMix (Roche) on a Bio-Rad C1000 Touch Thermocycler, using primer-specific thermal profiles. Products were resolved on 1-2% agarose gels in 1× TAE, stained with GelRed (Sigma-Aldrich), purified using the QIAquick PCR Purification Kit (QIAGEN), and submitted for Sanger sequencing (LGC Genomics) to confirm precise junction sequences. Digital PCR (dPCR) for Quantification: To quantify knock-in efficiency and transgene copy number, we performed droplet-free digital PCR on the QIAcuity platform (QIAGEN). Duplex probe-based assays measured transgene abundance relative to the single-copy reference gene AFF3. Genomic DNA was digested with XbaI (0.05 U/μL, NEB) to reduce viscosity, then partitioned into QIAcuity Nanoplates (12 μL for 96-well or 40 μL for 24-well format) containing QIAcuity Probe PCR Master Mix (1×), 800 nM primers for the transgene, 400 nM primers for AFF3, 400 nM FAM-labeled transgene probe, 200 nM HEX-labeled AFF3 probe, and 2-10 ng of digested DNA. For internal-internal junction assays, cycling conditions included enzyme activation at 95°C for 2 min, 40 cycles of 95°C for 15 s and 58°C for 30 s, followed by imaging. Outward-inward assays used 55 cycles and a final 2-min extension at 72°C. Absolute copy numbers were calculated by QIAcuity software using Poisson statistics and normalized to AFF3, expressed as copies per 100 cells. No-template controls were included on each plate, and fluorescence thresholds were manually reviewed to confirm assay specificity. Collectively, these three orthogonal methods provided complementary data on transgene expression, cell viability, enrichment of edited populations, and precise verification of genomic integration.

#### Assessment of Proliferation and Viability in CRISPR-Edited Neuroblastoma Lines

To compare growth kinetics and survival between CRISPR/Cas9-engineered and parental neuroblastoma cells, we used SK-N-BE(2)-C and SK-N-AS clones carrying targeted integrations alongside their wild-type counterparts. For each line, 1 × 10^4^ cells per well were dispensed into 96-well flat-bottom plates in biological duplicate and technical triplicate. Plates were transferred to the IncuCyte live-cell imaging system (Sartorius), which captured phase-contrast images every 2-4 hours over a 7-day period. These images were analyzed by the IncuCyte software to produce real-time proliferation curves. Parallel wells were supplemented at seeding with 250 nM Cytotox Green reagent (Sartorius, Cat. No. 4633) to label membrane-compromised cells. Green fluorescence was recorded alongside phase-contrast images, and the percentage of nonviable cells was calculated at each time point. This dual imaging approach enabled continuous, noninvasive tracking of both cell growth and viability in edited versus control populations.

#### Boyden Chamber Transwell Migration Assays

The functional activity of CXCL10 was evaluated employing a 24-well Boyden chamber transwell system equipped with polycarbonate membranes featuring 8 μm pores. Prior to cell seeding, transwell inserts were coated with 0.5% bovine serum albumin (BSA) in PBS for 1 hour at room temperature. T cells, isolated from donor peripheral blood mononuclear cells (PBMCs) using MACS columns and the Pan T Cell Isolation Kit (Miltenyi Biotec), were subsequently activated with CD3/CD28 Dynabeads (Thermo Fisher Scientific) before being resuspended at 5 × 10⁶ cells/mL for seeding into the upper transwell chamber. The lower chamber was filled with 500 μL of either conditioned medium derived from HEK293T cells expressing cytokines or corresponding control medium. Cell migration was recorded over a period of 4 hours using the IncuCyte S3 live-cell imaging platform, and migration data were subsequently analyzed with the associated IncuCyte S3 software.

#### Cytokine Quantification by ELISA

Secreted cytokines were measured by sandwich ELISA according to each manufacturer’s instructions. Human CXCL10/IP-10 and CXCL11/I-TAC levels were determined using DuoSet kits (R&D Systems, Cat. Nos. DY266 and DY392). High-binding 96-well plates (NUNC) were coated overnight at 4 °C with the appropriate capture antibody. After three washes in PBS-Tween (0.05%), wells were blocked for one hour at room temperature with 1% bovine serum. Thawed supernatants and standards were added in duplicate at dilutions of 1:10, 1:100, or undiluted to ensure readings fell within the linear range of the assay. Plates were incubated for two hours at room temperature, washed, and then incubated with biotinylated detection antibodies for one hour. Following additional washes, streptavidin-horseradish peroxidase was added for 30 minutes, and color was developed with TMB substrate for 20 minutes in the dark. The reaction was stopped with 2 N sulfuric acid (50 µL per well), and absorbance was read at 450 nm (570 nm reference) on an Epoch plate reader (Gen5 software). Concentrations were calculated from standard curves generated by four-parameter logistic regression.

#### Hematoxylin and Eosin (H&E) Staining

H&E staining was performed on paraffin-embedded tumor tissues. Tumors were excised, fixed in 4% neutral-buffered formalin at room temperature for 24-48 hours, dehydrated through a graded ethanol series, cleared in xylene, and embedded in paraffin. Sections were cut into 6 µm slices using a microtome and mounted onto positively charged glass slides. Slides were deparaffinized in xylene twice for 5 minutes each, followed by rehydration through a descending ethanol series (100%, 95%, 70%, and 50% ethanol, 5 minutes each) and rinsed in distilled water for 2 minutes. Sections were stained in Mayer’s hematoxylin solution (Sigma-Aldrich) for 5 minutes to visualize nuclei, then rinsed in running tap water for 5 minutes to remove excess stain. Differentiation was performed in 0.3% acid alcohol (1% HCl in 70% ethanol) for a few seconds until background cleared, followed by another rinse in tap water for 5 minutes. Sections were then blued in Scott’s tap water substitute (or 0.1% sodium bicarbonate solution) for 1 minute and rinsed again in tap water. Slides were counterstained in eosin Y solution (Sigma-Aldrich) for 2 minutes to stain cytoplasm and extracellular matrix components, dehydrated through ascending ethanol series (70%, 95%, 100% ethanol, 1 minute each), cleared in xylene twice for 2 minutes each, and mounted with a xylene-based mounting medium (e.g., DPX). Slides were air-dried and imaged using a bright-field microscope.

### Supplementary Notes

#### Supplementary Note 1: Reporter Benchmarking Reveals Cell Line–Specific GFP, RFP, and Q8 Expression Patterns

Correlation analyses between transfection efficiency and reporter accumulation revealed cell line-specific patterns **(Supplementary Figure S2b).** In SK-N-AS, GFP showed a strong positive correlation between reporter-positive percentages and MFI (p < 0.0001), while RFP showed no significant correlation, suggesting differences in translation or stability. In contrast, SK-N-BE(2)-C cells exhibited significant positive correlations for both GFP (p < 0.0001) and RFP (p < 0.001), indicating a robust link between transfection rates and per-cell **expression (Supplementary Figure S2b).** Comparative testing of monocistronic and bicistronic vectors showed RFP constructs yielded higher percentages of positive cells than GFP constructs, whereas Q8 surface-antigen expression was comparable to RFP in both cell lines **(Figure 1d).** These cell line–specific reporter patterns likely reflect both reporter properties and intrinsic biological differences. In SK-N-AS, heterogeneous plasmid uptake and slower proliferation lead to a small subset of highly expressing cells, resulting in low GFP⁺ percentages but disproportionately high GFP MFI. Differences in reporter maturation and turnover further uncouple the relationship between % positivity and MFI, explaining the weak RFP-MFI correlation in this line. In contrast, SK-N-BE(2)-C cells show more uniform transfection and reporter turnover, producing strong GFP and RFP correlations and proportional scaling of MFI with transfection rate. These findings highlight that reporter-based readouts behave nonlinearly in solid tumor models and must be interpreted in a cell-line-specific manner.

#### ****Supplementary Note 2: Electroporation Platform Benchmarking and Enhancer Testing****

Comparative platform analysis showed that the Lonza 4D-Nucleofector produced higher initial knock-in rates than the Neon Transfection System but was associated with slightly increased cytotoxicity **(Supplementary Figure S6a,b)**. Longitudinal expression analysis demonstrated that smaller constructs (e.g., CMVp_RFP_bGH) maintained stable expression for more than 35 days post electroporation across both platforms, whereas larger constructs (e.g., CMVp_CXCL10-P2A-RFP_sPA) exhibited higher long-term stability when delivered via the 4D-Nucleofector **(Supplementary Figure S6c)**. Testing IDT Enhancer v2, a DNA-PK inhibitor reported to increase HDR efficiency in other primary and cancer cell types, did not improve knock-in rates in either neuroblastoma cell line investigated **(Supplementary Figure S6d).**

#### ****Supplementary Note 3: ApoE Isoform Effects on LNP Uptake and Expression****

Comparison of ApoE isoforms revealed that both ApoE3 and ApoE4 enhanced LNP-mediated uptake and GFP mRNA transfection efficiency in neuroblastoma cells. However, SK-N-BE(2)-C cells showed higher GFP mRNA levels when supplemented with ApoE4 compared to ApoE3 (Supplementary Figure S9b). This suggests cell line-specific LNP. ApoE interactions, potentially reflecting differences in LDLR-family receptor engagement consistent with metabolic distinctions between adrenergic and mesenchymal neuroblastoma states **(Supplementary Figure S9b)**.

### Supplementary Figures

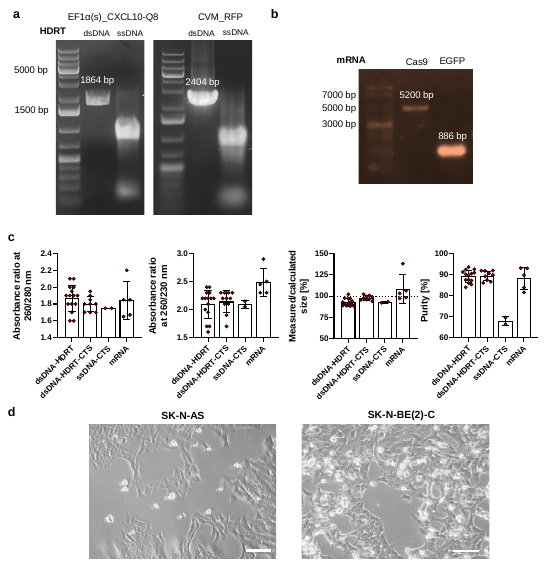

Supplementary Figure S1. Quality control of genome editing components and morphological characteristics of neuroblastoma cell lines. **(a)** Gel electrophoresis verifying HDR template sizes. **(b)** Gel analysis of Cas9 and EGFP mRNAs. **(c)** Spectrophotometric quality metrics across DNA and mRNA samples. **(d)** Microscopy images showing distinct cell morphologies between SK-N-AS and SK-N-BE(2)-C. *Data presentation: Means ± SD where applicable.*

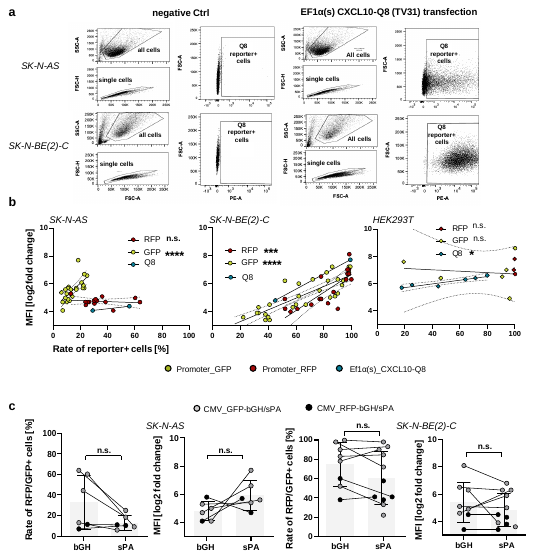

Supplementary Figure S2. Flow cytometric analysis of transgene expression for modular vector expression testing in neuroblastoma cells. **(a)** Gating strategy for detecting reporter-positive cells using PE-conjugated CD34 antibody (Qbend) for the Q8 epitope. **(b)** Correlations between reporter-positive percentages and MFI across different reporters and cell lines. **(c)** Comparison of reporter expression in constructs with bGH vs. synthetic polyadenylation signals*. Data presentation: Means ± SD. Statistical analysis: (b) Two-tailed Spearman correlation; (c) Mixed-effects model; p values: *<0.05, ***<0.001, ****<0.0001; n.s., not significant.*

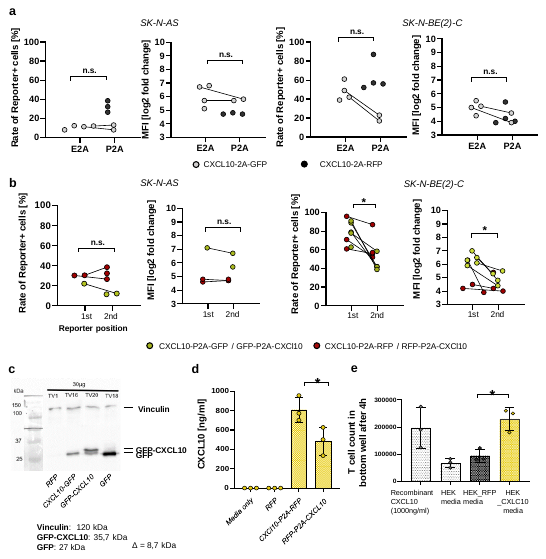

Supplementary Figure S3. Analysis of self-cleaving peptide efficiency, reporter gene positioning, and functional validation of secreted CXCL10. **(a)** Comparison of E2A vs. P2A linkers for GFP and RFP expression. **(b)** Impact of reporter gene positioning in bicistronic constructs on expression levels. **(c)** Western blot showing cleavage efficiency of mono- and bicistronic constructs. **(d)** ELISA quantification of CXCL10 secretion. **(e)** Transwell migration assay confirming biological activity, i.e. chemotactic function of secreted CXCL10. *Data presentation: Means ± SD. Statistical analysis: (a,b) matched two-way ANOVA, (d,e) Wilcoxon matched-pairs signed-rank test with Bonferroni correction; p values: *<0.05; n.s., not significant.*

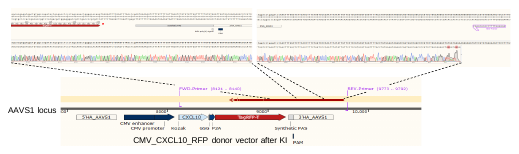

Supplementary Figure S4. Knock-in validation. Sanger sequencing confirming precise integration at AAVS1 locus.

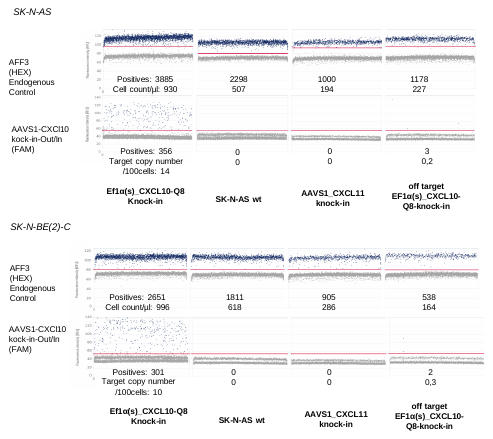

Supplementary Figure S5. Digital PCR validation of CRISPR-mediated knock-in events. dPCR scatter plots for AFF3 control and AAVS1 and control (off-target) knock-in of CXCL10 or CXCL11 using a junction spanning out-in assay specific for CXCl10 knock-in at the AAVS1 locus (FAM) and an AFF3 assay (HEX) as an endogenous control.

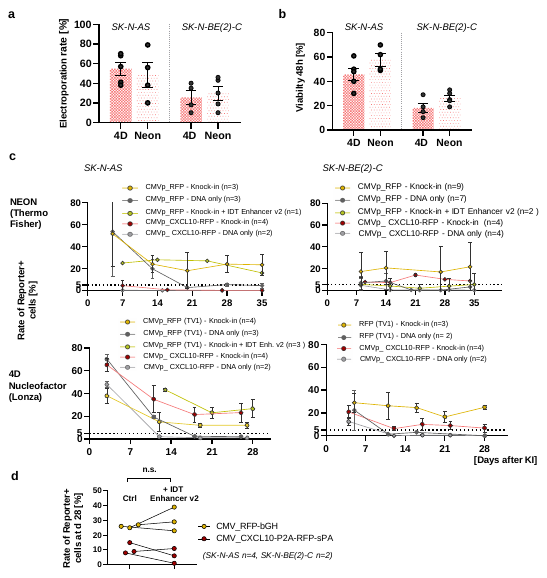

Supplementary Figure S6. Comparative performance of Neon vs. Lonza 4D electroporation platforms. **(a)** Electroporation efficiency across platforms. **(b)** Cell viability 48h post-electroporation. **(c)** Longitudinal transgene expression. **(d)** Knock-in rate 28d after electroporation measured as reporter positivity via flow cytometry with and without the use of IDT Enhancer v2. *Data presentation: Means ± SD. Statistical analysis: (d) Wilcoxon matched-pairs signed rank test; n.s., not significant.*

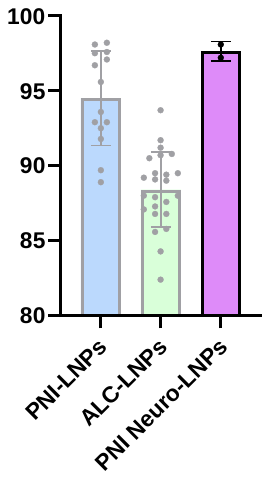

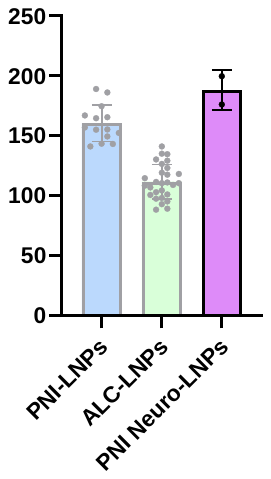

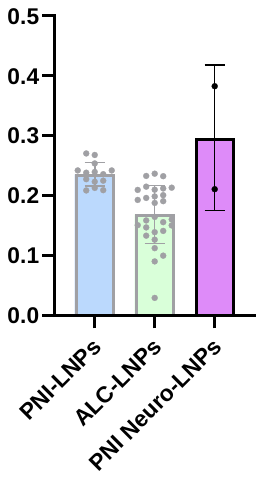

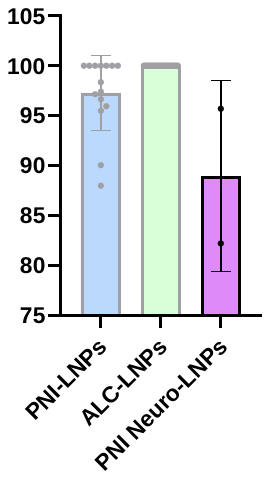

**size [nm]**

**PDI**

**Peak 1 area [%]**

**encapsulation efficacy [%]**

Supplementary Figure S7. Additional characterization of LNP formulations and cytotoxic effects. Dynamic light scattering (DLS) parameters (particle size, polydispersity index (PDI) and Peak 1 area for product purity) and mRNA encapsulation efficacy measured via ribo green assay for PNI Neuro-LNPs. *Data presentation: Means ± SD. Statistical analysis: (a) Mann-Whitney test. p values: **<0.01, ****<0.0001.*

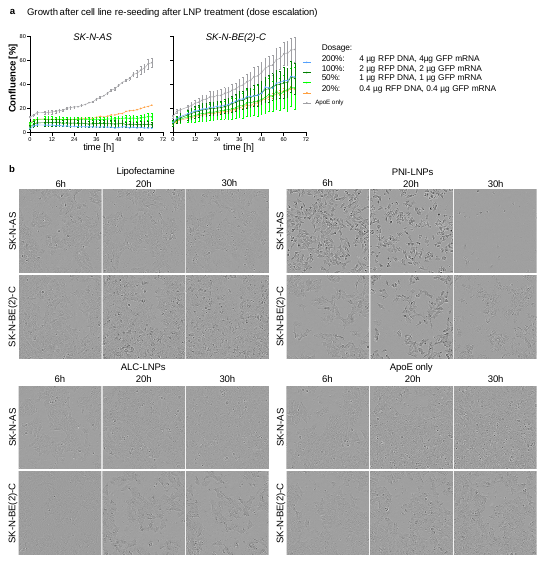

Supplementary Figure S8. LNP cytotoxic effects. **(a)** Growth changes with dose variation after re-seeding of the SK-N-AS and SK-N-BE(2)-C cell lines after LNP treatment with escalating doses. **(b)** IncuCyte imaging of morphological changes upon LNP treatment (100% dose from (a)). *Data presentation: Means ± SD.*

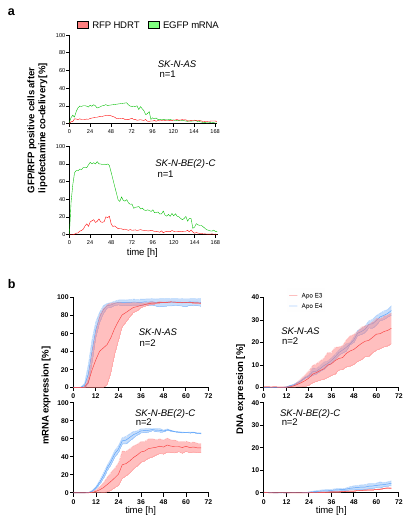

Supplementary Figure S9. Co-delivery of RFP DNA and GFP mRNA. **(a)** IncuCyte live-cell imaging data of RFP DNA and EGFP mRNA expression over 168 hours after Lipofectamine transfection in SK-N-AS and SK-N-BE(2)-C cell lines. **(b)** Nucleic acid expression detected via IncuCyte live-cell imaging after LNP transfection comparing Apolipoprotein E3 and E4.

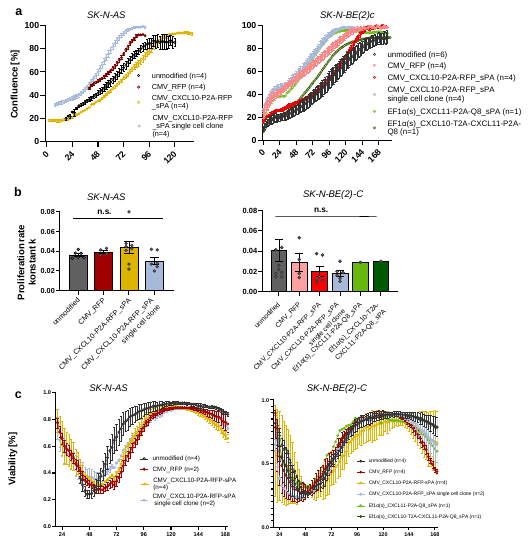

Supplementary Figure S10. Growth characterization of genetically modified neuroblastoma cell lines. **(a)** Confluence analysis over time. **(b)** Proliferation rates (growth constant k) as determined through logistic regression analysis of growth curves from (a). **(c)** Viability profiles in continuous culture measured via Incucyte live cell imaging. *Data presentation: Means ± SD. Statistical analysis: (b) One-Way-ANOVA with Dunn’s multiple comparisons test.*

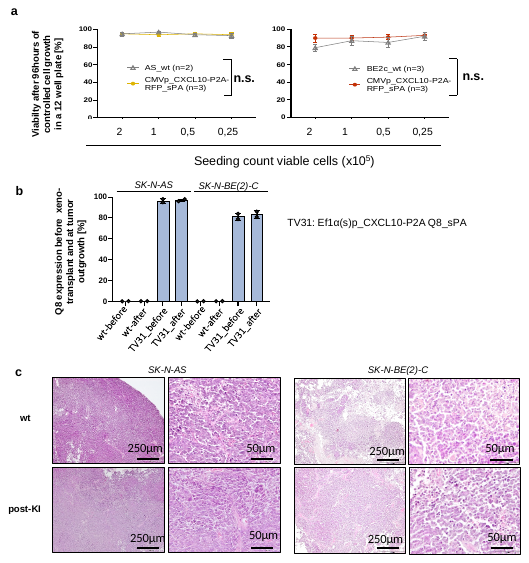

Supplementary Figure S11. Morphological and viability characterization of CRISPR-edited cell lines. **(a)** Viability at different seeding densities before subcutaneous injection of NOD mice. **(b)** Flow cytometric Q8 expression analysis before xeno-transplant and at tumor outgrowth. **(c)** H&E histology showing preserved neuroblastoma morphology post knock-in. *Data presentation: Means ± SD. Statistical analysis (a): TWO-WAY-ANOVA; n.s., not significant.*

### Supplementary Tables

#### Supplementary Table 1. List of transgene vectors

| **N°** | **Transgene vector** | **Temlpate size [bp]** | **N°** | **Transgene vector** | **template size [bp]** |
| --- | --- | --- | --- | --- | --- |
| TV1 | CMVp_RFP_bGH | 1604 | TV23 | MNDp_RFP_sPA | 1246 |
| TV2 | CMVp_CXCL10-E2A-EGFP_bGH | 2010 | TV24 | MNDp_GFP_sPA | 1248 |
| TV16 | CMVp_CXCL10-E2A-EGFP_sPA | 1814 | TV25 | Ef1α(s)p-EGFP_sPA | 1012 |
| TV17 | CMVp_CXCL10-P2A-RFP_sPA | 1759 | TV26 | CMVp_CXCL10-P2A-EGFP_sPA | 1783 |
| TV18 | CMVp_GFP_sPA | 1401 | TV28 | Ef1α_CXCL10-P2A-EGFP_sPA |  |
| TV19 | CMVp_RFP-P2A-CXCL10_sPA | 1770 | TV29 | Ef1α(s)p_CXCL10_sPA | 1759 |
| TV20 | CMVp_GFP-P2A-CXCL10_sPA | 1785 | TV30 | Ef1α(s)p_CXCL10-P2A-EGFP_sPA | 1374 |
| TV21 | Ef1αp_RFP_sPA | 1142 | TV31 | Ef1α(s)p_CXCL10-P2A Q8_sPA | 1065 |
| TV22 | EF1αp_GFP_sPA | 1147 | TV50 | Ef1α(s)p_CXCL11-P2A-Q8_sPA | 1147 |
|  |  |  | TV51 | Ef1α(s)p_CXCL10_T2A_CXCL11_P2A_Q8 | 1246 |

*bp: base pairs, bGH: Bovine Growth Hormone polyadenylation signal, CMV promoter: Cytomegalovirus Immediate-Early Promoter, CXCL10: C-X-C Motif Chemokine Ligand 10, CXCL11: C-X-C Motif Chemokine Ligand 11, E2A: Thosea asigna virus 2A peptide (self-cleaving peptide sequence), Ef1α promoter: Elongation Factor 1 Alpha Promoter, Ef1α(s) promoter: Custom designed shortened Elongation Factor 1 Alpha Promoter, GFP: Green Fluorescent Protein, HDRT: Homology Directed Repair Template, MND promoter: Myeloproliferative sarcoma virus enhancer, Negative control region deleted, DL587rev primer binding site, P2A: Porcine Teschovirus 2A peptide (self-cleaving peptide sequence), Q8 reporter tag: Custom reporter epitope tag (CD34 epitope and CD8 transmembrane domain), RFP: Red Fluorescent Protein, sPA: Short Polyadenylation Signal, T2A: Thosea asigna virus 2A peptide (self-cleaving peptide sequence)*

#### Supplementary Table 2. Sequences of elements used in HDRT constructs & mRNA

| **AAVS1 5’homology arm (399bp)** |
| --- |
| ACCGTTTTTCTGGACAACCCCAAAGTACCCCGTCTCCCTGGCTTTAGCCACCTCTCCATCCTCTTGCTTTCTTTGCCTGGACACCCCGTTCTCCTGTGGATTCGGGTCACCTCTCACTCCTTTCATTTGGGCAGCTCCCCTACCCCCCTTACCTCTCTAGTCTGTGCTAGCTCTTCCAGCCCCCTGTCATGGCATCTTCCAGGGGTCCGAGAGCTCAGCTAGTCTTCTTCCTCCAACCCGGGCCCCTATGTCCACTTCAGGACAGCATGTTTGCTGCCTCCAGGGATCCTGTGTCCCCGAGCTGGGACCACCTTATATTCCCAGGGCCGGTTAATGTGGCTCTGGTTCTGGGTACTTTTATCTGTCCCCTCCACCCCACAGTGGGGCCACTAGGGACAG |
| **AAVS1 3’homology arm (400bp)** |
| GATTGGTGACAGAAAAGCCCCATCCTTAGGCCTCCTCCTTCCTAGTCTCCTGATATTGGGTCTAACCCCCACCTCCTGTTAGGCAGATTCCTTATCTGGTGACACACCCCCATTTCCTGGAGCCATCTCTCTCCTTGCCAGAACCTCTAAGGTTTGCTTACGATGGAGCCAGAGAGGATCCTGGGAGGGAGAGCTTGGCAGGGGGTGGGAGGGAAGGGGGGGATGCGTGACCTGCCCGGTTCTCAGTGGCCACCCTGCGCTACCCTCTCCCAGAACCTGAGCTGCTCTGACGCGGCCGTCTGGTGCGTTTCACTGATCCTGGTGCTGCAGCTTCCTTACACTTCCCAAGAGGAGAAGCAGTTTGGAAAAACAAAATCAGAATAAGTTGGTCCTGAGTTCT |
| **bGH (206bp)** |
| CTGTGCCTTCTAGTTGCCAGCCATCTGTTGTTTGCCCCTCCCCCGTGCCTTCCTTGACCCTGGAAGGTGCCACTCCCACTGTCCTTTCCTAATAAAATGAGGAAATTGCATCGCATTGTCTGAGTAGGTGTCATTCTATTCTGGGGGGTGGGGTGGGGCAGGACAGCAAGGGGGAGGATTGGGAAGAgAATAGCAGGCATGCTGGG |
| **CMV promoter (508bp)** |
| CGTTACATAACTTACGGTAAATGGCCCGCCTGGCTGACCGCCCAACGACCCCCGCCCATTGACGTCAATAATGACGTATGTTCCCATAGTAACGCCAATAGGGACTTTCCATTGACGTCAATGGGTGGAGTATTTACGGTAAACTGCCCACTTGGCAGTACATCAAGTGTATCATATGCCAAGTACGCCCCCTATTGACGTCAATGACGGTAAATGGCCCGCCTGGCATTATGCCCAGTACATGACCTTATGGGACTTTCCTACTTGGCAGTACATCTACGTATTAGTCATCGCTATTACCATGGTGATGCGGTTTTGGCAGTACATCAATGGGCGTGGATAGCGGTTTGACTCACGGGGATTTCCAAGTCTCCACCCCATTGACGTCAATGGGAGTTTGTTTTGGCACCAAAATCAACGGGACTTTCCAAAATGTCGTAACAACTCCGCCCCATTGACGCAAATGGGCGGTAGGCGTGTACGGTGGGAGGTCTATATAAGCAGAGCT |
| **CXCL10 (294bp)** |
| ATGAACCAGACCGCCATCCTGATCTGCTGCCTGATCTTCCTGACCCTGAGCGGCATCCAGGGAGTCCCCCTGTCCAGAACAGTGCGGTGCACCTGTATCAGCATCTCTAATCAACCTGTGAACCCAAGAAGCCTGGAAAAGCTCGAGATCATCCCTGCTTCTCAGTTCTGCCCTAGAGTGGAAATCATCGCCACAATGAAAAAGAAGGGCGAGAAGAGATGTCTGAACCCCGAGAGCAAGGCCATTAAGAACCTGCTGAAGGCCGTGTCCAAAGAGATGAGCAAGCGGAGCCCT |
| **CXCL11 (282bp)** |
| ATGAGTGTGAAGGGCATGGCTATAGCCTTGGCTGTGATATTGTGTGCTACAGTTGTTCAAGGCTTCCCCATGTTCAAAAGAGGACGCTGTCTTTGCATAGGCCCTGGGGTAAAAGCAGTGAAAGTGGCAGATATTGAGAAAGCCTCCATAATGTACCCAAGTAACAACTGTGACAAAATAGAAGTGATTATTACCCTGAAAGAAAATAAAGGACAACGATGCCTAAATCCCAAATCGAAGCAAGCAAGGCTTATAATCAAAAAAGTTGAAAGAAAGAATTTT |
| **E2A (69bp)** |
| GGCAGTGGACAGTGTACTAATTATGCTCTCTTGAAATTGGCTGGAGATGTTGAGAGCAACCCTGGACCT |
| **Ef1α promoter (330bp)** |
| GTTTATTACAGGGACAGCAGAGATCCAGTTTGGGGATCAATTGCATGAAGAATCTGCTTAGGGTTAGGCGTTTTGCGCTGCTTCGCGAGGATCTGCGATCGCTCCGGTGCCCGTCAGTGGGCAGAGCGCACATCGCCCACAGTCCCCGAGAAGTTGGGGGGAGGGGTCGGCAATTGAACCGGTGCCTAGAGAAGGTGGCGCGGGGTAAACTGGGAAAGTGATGTCGTGTACTGGCTCCGCCTTTTTCCCGAGGGTGGGGGAGAACCGTATATAAGTGCAGTAGTCGCCGTGAACGTTCTTTTTCGCAACGGGTTTGCCGCCAGAACACAG |
| **Ef1α(s) promoter (239bp)** |
| GCTCCGGTGCCCGTCAGTGGGCAGAGCGCACATCGCCCACAGTCCCCGAGAAGTTGGGGGGAGGGGTCGGCAATTGAACCGGTGCCTAGAGAAGGTGGCGCGGGGTAAACTGGGAAAGTGATGTCGTGTACTGGCTCCGCCTTTTTCCCGAGGGTGGGGGAGAACCGTATATAAGTGCAGTAGTCGCCGTGAACGTTCTTTTTCGCAACGGGTTTGCCGCCAGAACACAGGCCGCCACC |
| **GFP (714bp)** |
| GTGAGCAAGGGCGAGGAGCTGTTCACCGGGGTGGTGCCCATCCTGGTCGAGCTGGACGGCGACGTAAACGGCCACAAGTTCAGCGTGTCCGGCGAGGGCGAGGGCGATGCCACCTACGGCAAGCTGACCCTGAAGTTCATCTGCACCACCGGCAAGCTGCCCGTGCCCTGGCCCACCCTCGTGACCACCCTGACCTACGGCGTGCAGTGCTTCAGCCGCTACCCCGACCACATGAAGCAGCACGACTTCTTCAAGTCCGCCATGCCCGAAGGCTACGTCCAGGAGCGCACCATCTTCTTCAAGGACGACGGCAACTACAAGACCCGCGCCGAGGTGAAGTTCGAGGGCGACACCCTGGTGAACCGCATCGAGCTGAAGGGCATCGACTTCAAGGAGGACGGCAACATCCTGGGGCACAAGCTGGAGTACAACTACAACAGCCACAACGTCTATATCATGGCCGACAAGCAGAAGAACGGCATCAAGGTGAACTTCAAGATCCGCCACAACATCGAGGACGGCAGCGTGCAGCTCGCCGACCACTACCAGCAGAACACCCCCATCGGCGACGGCCCCGTGCTGCTGCCCGACAACCACTACCTGAGCACCCAGTCCGCCCTGAGCAAAGACCCCAACGAGAAGCGCGATCACATGGTCCTGCTGGAGTTCGTGACCGCCGCCGGGATCACTCTCGGCATGGACGAGCTGTACAAG |
| **MND promoter (430bp)** |
| TCGGCAGCGGATTTCTAGAACTAGTATTAAGGATCCGAACAGAGAGACAGCAGAATATGGGCCAAACAGGATATCTGTGGTAAGCAGTTCCTGCCCCGCTCAGGGCCAAGAACAGTTGGAACAGCAGAATATGGGCCAAACAGGATATCTGTGGTAAGCAGTTCCTGCCCCGCTCAGGGCCAAGAACAGATGGTCCCCAGATGCGGTCCCGCCCTCAGCAGTTTCTAGAGAACCATCAGATGTTTCCAGGGTGCCCCAAGGACCTGAAATGACCCTGTGCCTTATTTGAACTAACCAATCAGTTCGCTTCTCGCTTCTGTTCGCGCGCTTCTGCTCCCCGAGCTCTATATAAGCAGAGCTCGTTTAGTGAACCGTCAGATCGCCTGGAGACGCCATCCACGCTGTTTTGACCTCCATAGAAGACACCGAC |
| **P2A (66bp)** |
| GGCAGTGGACAGTGTACTAATTATGCTCTCTTGAAATTGGCTGGAGATGTTGAGAGCAACCCTGGACCT |
| **Q8 reporter tag (417bp)** |
| ATGGGATTGGTTCGCCGAGGAGCGCGAGCAGGTCCCCGGATGCCCCGCGGATGGACTGCCCTGTGCCTGCTTTCCCTCCTTCCAAGCGGCTTCATGGCAGAATTGCCAACTCAAGGTACTTTCTCAAATGTTTCCACGAACGTTAGTCCAGCTAAACCCACTACAACACCTGCACCTAGACCCCCGACGCCAGCCCCTACTATAGCTTCTCAACCACTGAGTCTTCGGCCAGAGGCATGCAGGCCCGCAGCAGGTGGGGCAGTGCACACAAGGGGTCTCGATTTTGCTTGTGACATTTATATTTGGGCTCCTTTGGCAGGGACCTGTGGGGTACTTCTGCTGAGTCTGGTGATCACGTTGTATTGTAACCACAGGAATAGGCGGCGGGTTTGTAAATGTCCACGACCGGTTGTCTGA |
| **RFP (705bp)** |
| ATGGTGTCTAAGGGCGAAGAGCTGATTAAGGAGAACATGCACATGAAGCTGTACATGGAGGGCACCGTGAACAACCACCACTTCAAGTGCACATCCGAGGGCGAAGGCAAGCCCTACGAGGGCACCCAGACCATGAGAATCAAGGTGGTCGAGGGCGGCCCTCTCCCCTTCGCCTTCGACATCCTGGCTACCAGCTTCATGTACGGCAGCAGAACCTTCATCAACCACACCCAGGGCATCCCCGATTTCTTTAAGCAGTCCTTCCCTGAGGGCTTCACATGGGAGAGAGTCACCACATACGAAGACGGGGGCGTGCTGACCGCTACCCAGGACACCAGCCTCCAGGACGGCTGCCTCATCTACAACGTCAAGATCAGAGGGGTGAACTTCCCATCCAACGGCCCTGTGATGCAGAAGAAAACACTCGGCTGGGAGGCCAACACCGAGACCCTGTACCCCGCTGACGGCGGCCTGGAAGGCAGAACCGACATGGCCCTGAAGCTCGTGGGCGGGGGCCACCTGATCTGCAACTTCAAGACCACATACAGATCCAAGAAACCCGCTAAGAACCTCAAGATGCCCGGCGTCTACTATGTGGACCACAGACTGGAAAGAATCAAGGAGGCCGACAAAGAGACCTACGTCGAGCAGCACGAGGTGGCTGTGGCCAGATACTGCGACCTCCCTAGCAAACTGGGGCACAAA |
| **sPA (49bp)** |
| AATAAAAGATCTTTATTTTCATTAGATCTGTGTGTTGGTTTTTTGTGTG |
| **T2A (63bp)** |
| GGCAGTGGAGAGGGCAGAGGAAGTCTGCTAACATGCGGTGACGTCGAGGAGAATCCTGGCCCA |
| **pBR322_5ÚTR_Cas9-2A-mCherry_3ÚTR (mRNA template)** |

*AAVS1: Adeno-Associated Virus Integration Site 1, bp: base pairs, bGH: Bovine Growth Hormone polyadenylation signal, CMV promoter: Cytomegalovirus Immediate-Early Promoter, CXCL10: C-X-C Motif Chemokine Ligand 10, CXCL11: C-X-C Motif Chemokine Ligand 11, E2A: Thosea asigna virus 2A peptide (self-cleaving peptide sequence), Ef1α promoter: Elongation Factor 1 Alpha Promoter, Ef1α(s) promoter: Custom designed shortened Elongation Factor 1 Alpha Promoter, GFP: Green Fluorescent Protein, HDRT: Homology Directed Repair Template, MND promoter: Myeloproliferative sarcoma virus enhancer, Negative control region deleted, DL587rev primer binding site, P2A: Porcine Teschovirus 2A peptide (self-cleaving peptide sequence), Q8 reporter tag: Custom reporter epitope tag (CD34 epitope and CD8 transmembrane domain), RFP: Red Fluorescent Protein, sPA: Short Polyadenylation Signal, T2A: Thosea asigna virus 2A peptide (self-cleaving peptide sequence)*

#### Supplementary Table 3. Primer and probe sequences

| **HDRT amplification primers** | **Orientation** | **Sequence** |
| --- | --- | --- |
| AAVS1_5’HA_FWD | FWD | ACCGTTTTTCTGGACAACCCC |
| AAVS1_3’HA_REV | REV | AGAACTCAGGACCAACTTATTCTGATTTTGT |
| **PAM frequency/ CRISPR cutting (out/out)** | **Orientation** | **Sequence** |
| AAVS1_PAM freq. /Cutting_FWD | FWD | GGCATCTCTCCTCCCTCA |
| AAVS1_PAM freq./Cutting_REV | REV | TGGGGACTAGAAAGGTGAAG |
| **dPCR Primers and probes: Transgene CNV (In/In) Assays** | | |
| CXCL10_Transgene_FWD | FWD | GGAAAAGCTCGAGATCATCC |
| CXCL10_Transgene_REV | REV | TCTTTTTCATTGTGGCGATGATT |
| CXCL10_Transgene_Probe | Probe | 6FAM-CTGCTTCTCAGTTCTGCCCTAGAGTGG-BHQ1 |
| **dPCR Primers and probes: Site-specific knock-in (Out/In) Assays** | | |
| Ef1a_ Out/In _REV | REV | GCACTTATATACGGTTCTCC |
| AAVS1_Out/In_FWD | FWD | GCCGTCTTCACTCGCTG |
| AAVS1_Out/In _Probe | Probe | 6FAM-TCCCTTGCGTCCCGCCTCCCCTT-BHQ1 |
| **dPCR Primers and probes: Endogenous Control Assay** | | |
| AFF3 FWD | FWD | CACCTAGCATGTGTGGCATT |
| AFF3 REV | REV | GCAGATCCAGGTCGTTGAAG |
| AFF3 probe | Probe | HEX-AACAACTCTTTCTGTCCCCCT-BHQ1 |

*HDRT: Homology Directed Repair Template, FWD: forward, REV: reverse, freq.: Frequency, dPCR: digital PCR, BHQ1: Black Hole Quencher 1*

#### Supplementary Table 4. mRNA sequences

| **GFP (886 bp)** |
| --- |
| CGCGGCCGCTAATACGACTCACTATAAGGAAATAAGAGAGAAAAGAAGAGTAAGAAGAAATATAAGAGCCACCATGGTATCCAAGGGTGAAGAATTGTTCACTGGCGTAGTTCCTATCTTGGTGGAGCTCGACGGAGATGTCAATGGACACAAGTTTTCAGTCAGTGGCGAAGGTGAGGGTGATGCAACTTACGGGAAGCTTACACTGAAATTCATTTGTACTACTGGCAAGTTGCCGGTGCCATGGCCCACGCTGGTAACGACCCTTACCTACGGTGTCCAATGTTTTTCTCGATACCCCGATCATATGAAACAGCATGACTTTTTTAAATCTGCTATGCCCGAGGGATACGTACAGGAAAGGACTATATTTTTTAAGGACGATGGTAACTATAAGACCCGCGCCGAAGTCAAGTTTGAGGGAGATACTCTTGTTAATCGAATCGAGCTCAAGGGGATAGATTTTAAGGAGGATGGCAACATTCTCGGCCACAAGCTGGAGTACAATTACAATAGTCACAATGTTTACATAATGGCTGATAAGCAAAAAAACGGCATCAAAGTTAATTTTAAAATTAGGCATAATATAGAAGACGGTTCAGTTCAACTCGCCGACCACTACCAGCAGAATACCCCTATTGGCGATGGGCCGGTTCTCCTGCCTGACAACCATTACCTCTCAACACAGTCTGCTTTGAGTAAAGACCCTAACGAGAAGCGAGATCATATGGTCTTGCTCGAGTTCGTAACGGCCGCCGGTATAACCCTGGGCATGGATGAGCTCTACAAATAAGCTGCCTTCTGCGGGGCTTGCCTTCTGGCCATGCCCTTCTTCTCTCCCTTGCACCTGTACCTCTTGGTCTTTGAATAAAGCCTGAGTAGGAAGAAAAAAAAAAAAAAAAAAAAAAAAAAAAAAAAAAAAAAAAAAAAAAAAAAAAAAAAAAAAAAAAAAAAAAAAAAAAAAAAAAAAAAAAAAAAAAAAAAAAAAAAAAAAAAAAAAAAAAAA |
| **Cas9-mCherry (5212 bp)** |
| CGCGGCCGCTAATACGACTCACTATAAGGAAATAAGAGAGAAAAGAAGAGTAAGAAGAAATATAAGAGCCACCATGGACTATAAGGACCACGACGGAGACTACAAGGATCATGATATTGATTACAAAGACGATGACGATAAGATGGCCCCAAAGAAGAAGCGGAAGGTCGGTATCCACGGAGTCCCAGCAGCCGACAAGAAGTACAGCATCGGCCTGGACATCGGCACCAACTCTGTGGGCTGGGCCGTGATCACCGACGAGTACAAGGTGCCCAGCAAGAAATTCAAGGTGCTGGGCAACACCGACCGGCACAGCATCAAGAAGAACCTGATCGGAGCCCTGCTGTTCGACAGCGGCGAAACAGCCGAGGCCACCCGGCTGAAGAGAACCGCCAGAAGAAGATACACCAGACGGAAGAACCGGATCTGCTATCTGCAAGAGATCTTCAGCAACGAGATGGCCAAGGTGGACGACAGCTTCTTCCACAGACTGGAAGAGTCCTTCCTGGTGGAAGAGGATAAGAAGCACGAGCGGCACCCCATCTTCGGCAACATCGTGGACGAGGTGGCCTACCACGAGAAGTACCCCACCATCTACCACCTGAGAAAGAAACTGGTGGACAGCACCGACAAGGCCGACCTGCGGCTGATCTATCTGGCCCTGGCCCACATGATCAAGTTCCGGGGCCACTTCCTGATCGAGGGCGACCTGAACCCCGACAACAGCGACGTGGACAAGCTGTTCATCCAGCTGGTGCAGACCTACAACCAGCTGTTCGAGGAAAACCCCATCAACGCCAGCGGCGTGGACGCCAAGGCCATCCTGTCTGCCAGACTGAGCAAGAGCAGACGGCTGGAAAATCTGATCGCCCAGCTGCCCGGCGAGAAGAAGAATGGCCTGTTCGGAAACCTGATTGCCCTGAGCCTGGGCCTGACCCCCAACTTCAAGAGCAACTTCGACCTGGCCGAGGATGCCAAACTGCAGCTGAGCAAGGACACCTACGACGACGACCTGGACAACCTGCTGGCCCAGATCGGCGACCAGTACGCCGACCTGTTTCTGGCCGCCAAGAACCTGTCCGACGCCATCCTGCTGAGCGACATCCTGAGAGTGAACACCGAGATCACCAAGGCCCCCCTGAGCGCCTCTATGATCAAGAGATACGACGAGCACCACCAGGACCTGACCCTGCTGAAAGCTCTCGTGCGGCAGCAGCTGCCTGAGAAGTACAAAGAGATTTTCTTCGACCAGAGCAAGAACGGCTACGCCGGCTACATTGACGGCGGAGCCAGCCAGGAAGAGTTCTACAAGTTCATCAAGCCCATCCTGGAAAAGATGGACGGCACCGAGGAACTGCTCGTGAAGCTGAACAGAGAGGACCTGCTGCGGAAGCAGCGGACCTTCGACAACGGCAGCATCCCCCACCAGATCCACCTGGGAGAGCTGCACGCCATTCTGCGGCGGCAGGAAGATTTTTACCCATTCCTGAAGGACAACCGGGAAAAGATCGAGAAGATCCTGACCTTCCGCATCCCCTACTACGTGGGCCCTCTGGCCAGGGGAAACAGCAGATTCGCCTGGATGACCAGAAAGAGCGAGGAAACCATCACCCCCTGGAACTTCGAGGAAGTGGTGGACAAGGGCGCTTCCGCCCAGAGCTTCATCGAGCGGATGACCAACTTCGATAAGAACCTGCCCAACGAGAAGGTGCTGCCCAAGCACAGCCTGCTGTACGAGTACTTCACCGTGTATAACGAGCTGACCAAAGTGAAATACGTGACCGAGGGAATGAGAAAGCCCGCCTTCCTGAGCGGCGAGCAGAAAAAGGCCATCGTGGACCTGCTGTTCAAGACCAACCGGAAAGTGACCGTGAAGCAGCTGAAAGAGGACTACTTCAAGAAAATCGAGTGCTTCGACTCCGTGGAAATCTCCGGCGTGGAAGATCGGTTCAACGCCTCCCTGGGCACATACCACGATCTGCTGAAAATTATCAAGGACAAGGACTTCCTGGACAATGAGGAAAACGAGGACATTCTGGAAGATATCGTGCTGACCCTGACACTGTTTGAGGACAGAGAGATGATCGAGGAACGGCTGAAAACCTATGCCCACCTGTTCGACGACAAAGTGATGAAGCAGCTGAAGCGGCGGAGATACACCGGCTGGGGCAGGCTGAGCCGGAAGCTGATCAACGGCATCCGGGACAAGCAGTCCGGCAAGACAATCCTGGATTTCCTGAAGTCCGACGGCTTCGCCAACAGAAACTTCATGCAGCTGATCCACGACGACAGCCTGACCTTTAAAGAGGACATCCAGAAAGCCCAGGTGTCCGGCCAGGGCGATAGCCTGCACGAGCACATTGCCAATCTGGCCGGCAGCCCCGCCATTAAGAAGGGCATCCTGCAGACAGTGAAGGTGGTGGACGAGCTCGTGAAAGTGATGGGCCGGCACAAGCCCGAGAACATCGTGATCGAAATGGCCAGAGAGAACCAGACCACCCAGAAGGGACAGAAGAACAGCCGCGAGAGAATGAAGCGGATCGAAGAGGGCATCAAAGAGCTGGGCAGCCAGATCCTGAAAGAACACCCCGTGGAAAACACCCAGCTGCAGAACGAGAAGCTGTACCTGTACTACCTGCAGAATGGGCGGGATATGTACGTGGACCAGGAACTGGACATCAACCGGCTGTCCGACTACGATGTGGACCATATCGTGCCTCAGAGCTTTCTGAAGGACGACTCCATCGACAACAAGGTGCTGACCAGAAGCGACAAGAACCGGGGCAAGAGCGACAACGTGCCCTCCGAAGAGGTCGTGAAGAAGATGAAGAACTACTGGCGGCAGCTGCTGAACGCCAAGCTGATTACCCAGAGAAAGTTCGACAATCTGACCAAGGCCGAGAGAGGCGGCCTGAGCGAACTGGATAAGGCCGGCTTCATCAAGAGACAGCTGGTGGAAACCCGGCAGATCACAAAGCACGTGGCACAGATCCTGGACTCCCGGATGAACACTAAGTACGACGAGAATGACAAGCTGATCCGGGAAGTGAAAGTGATCACCCTGAAGTCCAAGCTGGTGTCCGATTTCCGGAAGGATTTCCAGTTTTACAAAGTGCGCGAGATCAACAACTACCACCACGCCCACGACGCCTACCTGAACGCCGTCGTGGGAACCGCCCTGATCAAAAAGTACCCTAAGCTGGAAAGCGAGTTCGTGTACGGCGACTACAAGGTGTACGACGTGCGGAAGATGATCGCCAAGAGCGAGCAGGAAATCGGCAAGGCTACCGCCAAGTACTTCTTCTACAGCAACATCATGAACTTTTTCAAGACCGAGATTACCCTGGCCAACGGCGAGATCCGGAAGCGGCCTCTGATCGAGACAAACGGCGAAACCGGGGAGATCGTGTGGGATAAGGGCCGGGATTTTGCCACCGTGCGGAAAGTGCTGAGCATGCCCCAAGTGAATATCGTGAAAAAGACCGAGGTGCAGACAGGCGGCTTCAGCAAAGAGTCTATCCTGCCCAAGAGGAACAGCGATAAGCTGATCGCCAGAAAGAAGGACTGGGACCCTAAGAAGTACGGCGGCTTCGACAGCCCCACCGTGGCCTATTCTGTGCTGGTGGTGGCCAAAGTGGAAAAGGGCAAGTCCAAGAAACTGAAGAGTGTGAAAGAGCTGCTGGGGATCACCATCATGGAAAGAAGCAGCTTCGAGAAGAATCCCATCGACTTTCTGGAAGCCAAGGGCTACAAAGAAGTGAAAAAGGACCTGATCATCAAGCTGCCTAAGTACTCCCTGTTCGAGCTGGAAAACGGCCGGAAGAGAATGCTGGCCTCTGCCGGCGAACTGCAGAAGGGAAACGAACTGGCCCTGCCCTCCAAATATGTGAACTTCCTGTACCTGGCCAGCCACTATGAGAAGCTGAAGGGCTCCCCCGAGGATAATGAGCAGAAACAGCTGTTTGTGGAACAGCACAAGCACTACCTGGACGAGATCATCGAGCAGATCAGCGAGTTCTCCAAGAGAGTGATCCTGGCCGACGCTAATCTGGACAAAGTGCTGTCCGCCTACAACAAGCACCGGGATAAGCCCATCAGAGAGCAGGCCGAGAATATCATCCACCTGTTTACCCTGACCAATCTGGGAGCCCCTGCCGCCTTCAAGTACTTTGACACCACCATCGACCGGAAGAGGTACACCAGCACCAAAGAGGTGCTGGACGCCACCCTGATCCACCAGAGCATCACCGGCCTGTACGAGACACGGATCGACCTGTCTCAGCTGGGAGGCGACAAAAGGCCGGCGGCCACGAAAAAGGCCGGCCAGGCAAAAAAGAAAAAGGAATTCGGCAGTGGAGAGGGCAGAGGAAGTCTGCTAACATGCGGTGACGTCGAGGAGAATCCTGGCCCAGTGAGCAAGGGCGAGGAGGATAACATGGCCATCATCAAGGAGTTCATGCGCTTCAAGGTGCACATGGAGGGCTCCGTGAACGGCCACGAGTTCGAGATCGAGGGCGAGGGCGAGGGCCGCCCCTACGAGGGCACCCAGACCGCCAAGCTGAAGGTGACCAAGGGTGGCCCCCTGCCCTTCGCCTGGGACATCCTGTCCCCTCAGTTCATGTACGGCTCCAAGGCCTACGTGAAGCACCCCGCCGACATCCCCGACTACTTGAAGCTGTCCTTCCCCGAGGGCTTCAAGTGGGAGCGCGTGATGAACTTCGAGGACGGCGGCGTGGTGACCGTGACCCAGGACTCCTCCCTGCAGGACGGCGAGTTCATCTACAAGGTGAAGCTGCGCGGCACCAACTTCCCCTCCGACGGCCCCGTAATGCAGAAGAAAACCATGGGCTGGGAGGCCTCCTCCGAGCGGATGTACCCCGAGGACGGCGCCCTGAAGGGCGAGATCAAGCAGAGGCTGAAGCTGAAGGACGGCGGCCACTACGACGCTGAGGTCAAGACCACCTACAAGGCCAAGAAGCCCGTGCAGCTGCCCGGCGCCTACAACGTCAACATCAAGTTGGACATCACCTCCCACAACGAGGACTACACCATCGTGGAACAGTACGAACGCGCCGAGGGCCGCCACTCCACCGGCGGCATGGACGAGCTGTACAAGTAAGCTGCCTTCTGCGGGGCTTGCCTTCTGGCCATGCCCTTCTTCTCTCCCTTGCACCTGTACCTCTTGGTCTTTGAATAAAGCCTGAGTAGGAAGAAAAAAAAAAAAAAAAAAAAAAAAAAAAAAAAAAAAAAAAAAAAAAAAAAAAAAAAAAAAAAAAAAAAAAAAAAAAAAAAAAAAAAAAAAAAAAAAAAAAAAAAAAAAAAAAAAAAAAAA |
| **Vector Backbone (pBR322)** |
| CCGGTCATCATCACCATCACCATTGAGTTTAAACCCGCTGATCAGCCTCGACTGTGCCTTCTAGTTGCCAGCCATCTGTTGTTTGCCCCTCCCCCGTGCCTTCCTTGACCCTGGAAGGTGCCACTCCCACTGTCCTTTCCTAATAAAATGAGGAAATTGCATCGCATTGTCTGAGTAGGTGTCATTCTATTCTGGGGGGTGGGGTGGGGCAGGACAGCAAGGGGGAGGATTGGGAAGACAATAGCAGGCATGCTGGGGATGCGGTGGGCTCTATGGCTTCTGAGGCGGAAAGAACCAGCTGGGGCTCGATACCGTCGACCTCTAGCTAGAGCTTGGCGTAATCATGGTCATAGCTGTTTCCTGTGTGAAATTGTTATCCGCTCACAATTCCACACAACATACGAGCCGGAAGCATAAAGTGTAAAGCCTAGGGTGCCTAATGAGTGAGCTAACTCACATTAATTGCGTTGCGCTCACTGCCCGCTTTCCAGTCGGGAAACCTGTCGTGCCAGCTGCATTAATGAATCGGCCAACGCGCGGGGAGAGGCGGTTTGCGTATTGGGCGCTCTTCCGCTTCCTCGCTCACTGACTCGCTGCGCTCGGTCGTTCGGCTGCGGCGAGCGGTATCAGCTCACTCAAAGGCGGTAATACGGTTATCCACAGAATCAGGGGATAACGCAGGAAAGAACATGTGAGCAAAAGGCCAGCAAAAGGCCAGGAACCGTAAAAAGGCCGCGTTGCTGGCGTTTTTCCATAGGCTCCGCCCCCCTGACGAGCATCACAAAAATCGACGCTCAAGTCAGAGGTGGCGAAACCCGACAGGACTATAAAGATACCAGGCGTTTCCCCCTGGAAGCTCCCTCGTGCGCTCTCCTGTTCCGACCCTGCCGCTTACCGGATACCTGTCCGCCTTTCTCCCTTCGGGAAGCGTGGCGCTTTCTCATAGCTCACGCTGTAGGTATCTCAGTTCGGTGTAGGTCGTTCGCTCCAAGCTGGGCTGTGTGCACGAACCCCCCGTTCAGCCCGACCGCTGCGCCTTATCCGGTAACTATCGTCTTGAGTCCAACCCGGTAAGACACGACTTATCGCCACTGGCAGCAGCCACTGGTAACAGGATTAGCAGAGCGAGGTATGTAGGCGGTGCTACAGAGTTCTTGAAGTGGTGGCCTAACTACGGCTACACTAGAAGAACAGTATTTGGTATCTGCGCTCTGCTGAAGCCAGTTACCTTCGGAAAAAGAGTTGGTAGCTCTTGATCCGGCAAACAAACCACCGCTGGTAGCGGTGGTTTTTTTGTTTGCAAGCAGCAGATTACGCGCAGAAAAAAAGGATCTCAAGAAGATCCTTTGATCTTTTCTACGGGGTCTGACGCTCAGTGGAACGAAAACTCACGTTAAGGGATTTTGGTCATGAGATTATCAAAAAGGATCTTCACCTAGATCCTTTTAAATTAAAAATGAAGTTTTAAATCAATCTAAAGTATATATGAGTAAACTTGGTCTGACAGTTACCAATGCTTAATCAGTGAGGCACCTATCTCAGCGATCTGTCTATTTCGTTCATCCATAGTTGCCTGACTCCCCGTCGTGTAGATAACTACGATACGGGAGGGCTTACCATCTGGCCCCAGTGCTGCAATGATACCGCGAGACCCACGCTCACCGGCTCCAGATTTATCAGCAATAAACCAGCCAGCCGGAAGGGCCGAGCGCAGAAGTGGTCCTGCAACTTTATCCGCCTCCATCCAGTCTATTAATTGTTGCCGGGAAGCTAGAGTAAGTAGTTCGCCAGTTAATAGTTTGCGCAACGTTGTTGCCATTGCTACAGGCATCGTGGTGTCACGCTCGTCGTTTGGTATGGCTTCATTCAGCTCCGGTTCCCAACGATCAAGGCGAGTTACATGATCCCCCATGTTGTGCAAAAAAGCGGTTAGCTCCTTCGGTCCTCCGATCGTTGTCAGAAGTAAGTTGGCCGCAGTGTTATCACTCATGGTTATGGCAGCACTGCATAATTCTCTTACTGTCATGCCATCCGTAAGATGCTTTTCTGTGACTGGTGAGTACTCAACCAAGTCATTCTGAGAATAGTGTATGCGGCGACCGAGTTGCTCTTGCCCGGCGTCAATACGGGATAATACCGCGCCACATAGCAGAACTTTAAAAGTGCTCATCATTGGAAAACGTTCTTCGGGGCGAAAACTCTCAAGGATCTTACCGCTGTTGAGATCCAGTTCGATGTAACCCACTCGTGCACCCAACTGATCTTCAGCATCTTTTACTTTCACCAGCGTTTCTGGGTGAGCAAAAACAGGAAGGCAAAATGCCGCAAAAAAGGGAATAAGGGCGACACGGAAATGTTGAATACTCATACTCTTCCTTTTTCAATATTATTGAAGCATTTATCAGGGTTATTGTCTCATGAGCGGATACATATTTGAATGTATTTAGAAAAATAAACAAATAGGGGTTCCGCGCACATTTCCCCGAAAAGTGCCACCTGACGTCGACGGATCGGGAGATCGATCTCCCGATCCCCTAGGGTCGACTCTCAGTACAATCTGCTCTGATGCCGCATAGTTAAGCCAGTATCTGCTCCCTGCTTGTGTGTTGGAGGTCGCTGAGTAGTGCGCGAGCAAAATTTAAGCTACAACAAGGCAAGGCTTGACCGACAATTGCATGAAGAATCTGCTTAGGGTTAGGCGTTTTGCGCTGCTTCGCGATGTACGGGCCAGATATACGCGTTGACATTGATTATTGACTAGTTATTAATAGTAATCAATTACGGGGTCATTAGTTCATAGCCCATATATGGAGTTCCGCGTTACATAACTTACGGTAAATGGCCCGCCTGGCTGACCGCCCAACGACCCCCGCCCATTGACGTCAATAATGACGTATGTTCCCATAGTAACGCCAATAGGGACTTTCCATTGACGTCAATGGGTGGAGTATTTACGGTAAACTGCCCACTTGGCAGTACATCAAGTGTATCATATGCCAAGTACGCCCCCTATTGACGTCAATGACGGTAAATGGCCCGCCTGGCATTATGCCCAGTACATGACCTTATGGGACTTTCCTACTTGGCAGTACATCTACGTATTAGTCATCGCTATTACCATGGTGATGCGGTTTTGGCAGTACATCAATGGGCGTGGATAGCGGTTTGACTCACGGGGATTTCCAAGTCTCCACCCCATTGACGTCAATGGGAGTTTGTTTTGGCACCAAAATCAACGGGACTTTCCAAAATGTCGTAACAACTCCGCCCCATTGACGCAAATGGGCGGTAGGCGTGTACGGTGGGAGGTCTATATAAGCAGAGCTGGTTTAGTGAACCGTCAGATCCGCTAGAGATC |

*The EGFP and Cas9_mCherry plasmids used as PCR templates for IVT were provided by the Wagner laboratory.*

#### Supplementary Table 5. Lipid mixes

| **Lipid Mix** | **Ionizable Lipid** | **Molar ratio** | **Phospholipid** | **Molar ratio** | **PEG Lipid** | **Molar ratio** | **Cholesterol** | | **Molar ratio** |
| --- | --- | --- | --- | --- | --- | --- | --- | --- | --- |
| **Lipid Mix A** | ALC-0315 | 50,0% | 18:0 DSPC | 10,0% | DMG-PEG (2000) | 2,0% | Cholesterol | | 38,0% |
| **Lipid Mix B** | ALC-0315 | 50,0% | 18:0 DSPC | 10,5% | DMG-PEG (2000)  DSPE-PEG (2000) Maleimid | 1,4%  0,1% | Cholesterol | | 38,0% |
| **Lipid Mix C** | ALC-0315 | 60,0% | 18:0 DSPC | 10,6% | DMG-PEG (2000) | 1,9% | Cholesterol | | 27,5% |

#### Supplementary Table 6. Equipment and Consumables used

| **Item** | **Catalogue n°** | **Provider** |
| --- | --- | --- |
| **Equipment** |  |  |
| 2100 Bioanalyzer Instrument |  | Agilent |
| C1000 Touch Thermal cycler | 1851148 | Bio-Rad |
| S220 Focused-ultrasonicator | 500217 | Covaris |
| Centrifuge 5418R |  | Eppendorf GmbH |
| Centrifuge 5424 |  | Eppendorf GmbH |
| Centrifuge 5427R |  | Eppendorf GmbH |
| Centrifuge 5810 R |  | Eppendorf GmbH |
| Duomax 1030, platform shaker |  | Heidolph instruments |
| Electronic Balance ABS 80-4 |  | Kern & Sohn GmbH |
| Electronic Rotary Microtome HM340E |  | Biotek |
| Erlenmeyer flasks |  | Thermo Scientific |
| FlowCytometer LSR-Fortessa X-20 |  | BD Biosciences |
| Fortessa Aria Cell sorter |  | BD Bioscience |
| Freezer (-20°C) LCv4010 |  | Carl Roth |
| Freezer (-80°C), HeraFreeze T series HFU400TV63 |  | Thermo Scientific |
| GloMaxR Multi |  | Promega |
| Heat controlled pressure cooker |  | Dako |
| Ice machine Manitowoc SOTTO |  | Manitowoc |
| Incubator HERAcell 240i CO2 |  | Thermo Scientific |
| Laminar Airflow Bench HERA safe 2020 KSP18 |  | Thermo Scientific |
| Luminometer GloMaxR- Multi+Microplate Multimode Reader with InstinctR E8032 |  | Promega |
| Magnetic Shaker RH basic |  | IKA Laboratoy Equipment |
| Manual system microscope Olympus BX43 |  | Olympus |
| Microscope Axio Vert.A1 |  | Zeiss |
| Millipore Barnstead MicroPure |  | Thermo Scientific |
| NanoDrop 2000 |  | Thermo Scientific |
| Neubauer counting chamber |  | Carl Roth |
| NextSeq 550 System |  | illumina |
| PCR workstation Pro |  | VWR Peqlab |
| pH-meter Five Easy Le409 |  | Mettler Toledo |
| Pipette filler pipetusR |  | Hirschmann Labortechnik |
| Pipette multipipetteR stream |  | Eppendorf GmbH |
| Pipettes (2.5 - 1000 μl) |  | Eppendorf GmbH |
| QuadroMACS |  | Miltenyi |
| Qubit fluorometer |  | Thermo Scientific |
| QIAcuity Digital PCR System (6-plex) |  | Qiagen |
| Shandon Excelsior ES |  | Thermo Scientific |
| Spectrophotometer EPOCH |  | BioTek Instruments |
| StepOnePlus Real-Time PCR system |  | Thermo Scientific |
| Suction pump AC02 |  | Carl Roth |
| Table centrifuge mini star silverline |  | VWR |
| ThermoMixer C |  | Eppendorf GmbH |
| Transwell Permeable supports |  | Costar |
| Ultracentrifuge Optima L90K |  | Beckman Coulter |
| Vortexer Reax top |  | Heidolph instruments |
| Vortexer VWRR Galaxy Mini Star |  | VWR International bvba |
| Waterbath GFL 1086 |  | GFL Technology |
| **General consumables** |  |  |
| 8-Well PCR tube strips plus 8 domed caps | strips: 72.985.002, caps: 65.989.002 | Sarstedt |
| 96-Well Plate Advanced TC (flat bottom) | 655983 | Greiner Bio-one |
| Biosphere® Filter Tip 10 µl | 70.1130.210 | Sarstedt |
| Biosphere® Filter Tip 100 µl | 70.760.212 | Sarstedt |
| Biosphere® Filter Tip 1000 µl | 70.762.211 | Sarstedt |
| Cell culture multiwell plate, 6 well, PS, clear, sterile | 657160 | Greiner Bio-one |
| Cell culture plates CELLSTAR®, sterile, white-96-well plates | KL43.1 | Roth |
| CELLSTAR®, TC, lid with condensation rings, sterile | 655180 | Greiner Bio-one |
| CryoPure Tube 1.6 ml yellow | 72.380.004 | Sarstedt |
| Disposable needles Sterican® long bevel facet, 0.30x12 mm | 4656300 | B.Braun |
| Disposable Syringe, Luer, 1 ml | CH030001L | Charina |
| FACS tubes | 352052 | BD |
| Falcon® 10 ml Serological Pipet | 357551 | Corning |
| Falcon® 12-well Clear Flat Bottom TC-treated Multiwell Cell Culture Plate | 353043 | Corning |
| Falcon® 15 ml High Clarity PP Centrifuge Tube | 352096 | Corning |
| Falcon® 2 ml Serological Pipet | 357507 | Corning |
| Falcon® 24-well Clear Flat Bottom TC-treated Multiwell Cell Culture Plate | 353047 | Corning |
| Falcon® 25 ml Serological Pipet | 357525 | Corning |
| Falcon® 35 mm TC-treated Easy-Grip Style Cell Culture Dish | 353001 | Corning |
| Falcon® 40 µm Cell Strainer | 352340 | Corning |
| Falcon® 48-well Clear Flat Bottom TC-treated Multiwell Cell Culture Plate | 353078 | Corning |
| Falcon® 5 ml Round Bottom Polystyrene Test Tube | 352052 | Corning |
| Falcon® 5 ml Round Bottom Polystyrene Test Tube, with Cell Strainer Snap Cap | 352235 | Corning |
| Falcon® 5 ml Serological Pipet | 357543 | Corning |
| Falcon® 50 ml High Clarity PP Centrifuge Tube | 352070 | Corning |
| FrameStar Fast Plate 96-well semi skirted | 4ti-1200 | 4titude |
| Injekt Solo-2-piece single-use syringe, 10 ml, Luer Lock | 201235 | B.Braun |
| MACS LS Columns | 130-042-401 | Miltenyi Biotec |
| MACS MS Columns | 130-042-201 | Miltenyi Biotec |
| Pasteurpipette glass, 145mm | 500635 | Brand |
| Pasteurpipette glass, 230mm | 500636 | Brand |
| Reagent Reservoirs, 25 ml | EKT8.1 | Carl Roth |
| Rotilab®-syringe filters, CA, sterile, 0.45µm | KC71.1 | Carl Roth |
| Sodium butyrate | B5887-1g | Sigma Aldrich |
| Spitzen Filter Surphob 100 µl (10x96) | VT0230 | Biozym |
| Spitzen Filter Surphob 10 µl lang (10x96) | VT0200 | Biozym |
| Spitzen Filter Surphob 1,250 µl (10x96) | VT0270 | Biozym |
| Stericup | CT92.1 | Carl Roth |
| Sterile, Stainless Steel, Premium Disposable Scalpel, 11PA | 03025 | Razormed |
| Transwell Permeable supports | 10107341 | Costar |
| Vasco® Nitril blue glove S | 9205518 | B.Braun |
| **Chemicals and reagents (general)** |  |  |
| 1-Step Ultra TMB ELISA Substrate | 34022 | Thermo Fischer Scientific |
| 0.05% Trypsin-EDTA (1x) | 25300-096 | Gibco |
| 2x HEPES | S0874 | Takara |
| Albumin from bovine serum Fraktion V | 8076.3 | Carl Roth |
| Anti-human CD28 antibody | 302934 | Biolegend |
| Anti-PE microbeads \| \| Invitrogen | 130-105-639 | Miltenyi |
| Brefeldin A | #347688 | BD Biosciences |
| CD3 antibody anti-human, pure-functional grade clone OKT3 | 130-093-387 | Miltenyi |
| CD8 microbeads, human - lyophilized | 130-097-057 | Miltenyi |
| CS&T Research Beads | 655050 | Becton Dickinson |
| Cut Smart Buffer | NEB#B6004 | New England Biolabs |
| D-Luciferin | 122799 | Perkin Elmer Inc. |
| Dimethyl sulfoxide (DMSO) | A994.1 | Carl Roth |
| DISPASE II | D4693-1G | Sigma Aldrich |
| Dnase I | A3778.0100 | AppliChem |
| Dynabeads Human T-Activator CD3/CD28 | 11131D | ThermoFisher |
| eosin Y solution | HT110116 | Sigma-Aldrich |
| EpiTect Hi-C Kit | 59971 | Qiagen |
| Ethanol Rotipuran 99.8% p.a. | 2065.2 | Carl Roth |
| Fetal Bovine Serum (FBS) SUPERIOR | S0615-500ML | Merck |
| Ficoll Paque Plus | 17144002 | Cytiva |
| Flow-Set Pro Fluorospheres | A62492 | Beckman Coulter |
| FlowCheck Beads | A63493 | Beckman Coulter |
| FlowClean Cleaning Agent | A64669 | Beckman Coulter |
| GelRed® | 12352106 | Sigma-Aldrich |
| Glycine Buffer ≥ 99% p.a. | 3908.3 | Carl Roth |
| H₂O₂ (2N) | 339741 | Sigma Aldrich |
| Human IL-15, premium grade | 130-095-765 | Miltenyi |
| Human IL-7, premium grade | 130-095-362 | Miltenyi |
| Human Fc Receptor Blocking Solution | 422301 | Biolegend |
| Hydrocortisone | H0396-100MG | Sigma Aldrich |
| Ionomycin | 10364-1MG | Sigma Aldrich |
| IsoFlow Sheath Fluid | 8546859 | Beckman Coulter |
| Isoflurane (CPS) | 1214 | Cp-pharma |
| KAPA HiFi HotStart Ready Mix | 7958927001 | Roche double ! |
| L-Glutamine | 25020-081 | Life Technologies |
| LE Agarose | 240004 | Biozym |
| Live/Dead Fixable Near IR (780) Viability Kit, for 633 nm excitation | L34975 | ThermoFisher Scientific |
| Matrigel | 356237 | Corning |
| Mayer’s hematoxylin solution | MHS80 | Sigma-Aldrich |
| Monensin | M5273-500MG | Sigma Aldrich |
| NextSeq 500/550 v2.5 Kits | Cat. Nummer fehlt | illumina |
| Nuclease-free water | 1097705 | Invitrogen |
| Papain from papaya latex buffered aqueous solution | P3128-100MG | Sigma Aldrich |
| Penicillin-Streptomycin (10,000 U/mL) | 18140122 | Gibco |
| PhosSTOP | 4906837001 | Sigma Aldrich |
| PMA | P8139-1MG | Sigma Aldrich |
| Potassium chloride (KCl) ≥ 99.5% p.a. | 6781.1 | Carl Roth |
| Potassium dihydrogen phosphate (KH₂PO₄) ≥ 99% p.a. | 3904.1 | Carl Roth |
| Potassium hydrogen carbonate (KHCO₃) ≥ 99% | X887.1 | Carl Roth |
| Powdered milk, blotting grade | T145.3 | Carl Roth |
| Precision Plus Protein Standards | 161-0374 | Bio-Rad |
| Proteinase K | 39450-01-6 | Carl Roth |
| Q5 Polymerase | M0491L | New England Biolabs |
| QIAquick Gel Extraction Kit | 28704 | Qiagen |
| QIAquick PCR Purification Kit | 28106 | Qiagen |
| QIAseq Library Quant Kit | 333314 | Qiagen |
| Retronectin | T1008 | Takara |
| SDS Pellets ≥ 99.9% | CN30.3 | Carl Roth |
| Streptavidin-Horseradish Peroxidase (HRP) Conjugate | SA10001 | Thermo Fischer Scientific |
| TEMED ≥ 98.5% | 2367.1 | Carl Roth |
| Tris Bufferan® ≥ 99.9% p.a. | 4855.5 | Carl Roth |
| Trypan Blue Solution 0.4% | 15250061 | Gibco |
| Tween®20 | 9127.2 | Carl Roth |
| Vibrant™ Dio Cell-Labeling Solution | V22886 | ThermoFisher |
| ZymoPURE Plasmid Miniprep Kit | D4211 | Zymo Research |
| β-Mercaptoethanol 99% p.a. | 4227 | Carl Roth |
| **Nanoparticle production** |  |  |
| NanoAssemblr Spark Cartridges - 20 Pack | NIS0009 | Preciscion NanoSystems |
| NanoAssemblr Spark Cartridges - 80 Pack | NIS0013 | Preciscion NanoSystems |
| GenVoy-ILM T Cell Kit for mRNA, SPARK | 1000683 | Preciscion NanoSystems |
| GenVoy-ILM - 2mL | NWW0041 | Preciscion NanoSystems |
| NanoAssemblr Spark Demo Kit - mRNA | NCS0017 | Preciscion NanoSystems |
| NanoAssemblr Spark Kit - Neuro9 siRNA - 2x2 nmol | NWS0001 | Preciscion NanoSystems |
| Spark Hepato 9 Kit -2x 2nmol | NWS0009 | Preciscion NanoSystems |
| PNI Formulation Buffer - 20mL | NWW0043 | Preciscion NanoSystems |
| ImmunoCultTM-XF T Cell Expansion Medium | 10981 | Stemcell Technologies Inc |
| Recombinant Human ApoE3 | 350-02 | Peprotech |
| Recombinant Human ApoE4 | 350-04 | Peprotech |
| **Lipids** |  |  |
| DLin-MC3-DMA (MC3) (50mg) | 25325 | Nanosoft polymers |
| DLin-MC3-DMA (MC3) (100mg) | 25325 | Nanosoft polymers |
| DLin-MC3-DMA (MC3) (1g) | 25325 | Nanosoft polymers |
| ALC-0315 | 890900O | Sigma-Aldrich / Merck (Avanti Polar Lipids) |
| Cholesterol | 700000P | Sigma-Aldrich (Avanti) |
| 18:0 DSPC (1,2-distearoyl-sn-glycero-3-phosphocholine) | 850365P | Sigma-Aldrich (Avanti) |
| 18:0 PEG2000 PE (1,2-distearoyl-sn-glycero-3-phosphoethanolamine-N-[methoxy(polyethylene glycol)-2000) | 880120P | Sigma-Aldrich (Avanti) |
| 14:0 PEG2000 PE (1,2-dimyristoyl-sn-glycero-3-phosphoethanolamine-N-[methoxy(polyethylene glycol)-2000] (ammonium salt)) | 880150P | Sigma-Aldrich (Avanti) |
| DMG-PEG (2000) (1,2-Dimyristoyl-rac-glycero-3-methoxypolyethylene glycol-2000) | 880151P | Sigma-Aldrich (Avanti) |
| DSPE-PEG (1,2-distearoyl-sn-glycero-3-phosphoethanolamine) | P3531 | Sigma-Aldrich (Avanti) |
| DSPE-PEG(2000) Carboxy-NHS | 880138P | Avanti |
| DSPE-PEG(2000) Maleimid (1,2-distearoyl-sn-glycero-3-phosphoethanolamine-N-[maleimide(polyethylene glycol)-2000) | 880126P | Avanti |
| Hydro Soy PC (L-α-phosphatidylcholine) | 840058P | Avanti |
| 18:1 TAP (DOTAP) 1,2-dioleoyl-3-trimethylammonium-propane | 890890P | Avanti |
| 18:1 (Δ9-Cis) PE (DOPE) (1,2-dioleoyl-sn-glycero-3-phosphoethanolamine) | 850725P | Avanti |
| **Tape station 4200 Agilent** |  |  |
| RNA ScreenTape (50bp-several kb) | 5067-5576 | Agilent |
| RNA ScreenTape Ladder | 5067-5578 | Agilent |
| RNA ScreenTape Sample Buffer | 5067-5577 | Agilent |
| D5000 ScreenTape (100 to 5000bp) | 5067-5588 | Agilent |
| D5000 Ladder | 5067-5590 | Agilent |
| D5000 Reagents | 5067-5589 | Agilent |
| **Cell lines** |  |  |
| SK-N-AS |  | Prof. Michael Claus V. Jensen |
| SK-N-BE(2)-C |  | Prof. Michael Claus V. Jensen |
| HEK293T | ATCC CRL-3216 | ATCC |
| **Cell culture media** |  |  |
| RPMI Medium 1640 (1x) | 21875-034 | Gibco™ |
| DMEM (1x) Dulbecco’s Modified Eagle Medium | 41966052 | Gibco™ |
| Endothelial Cell Growth Medium 2 | C-22111/39211 | PromoCell |
| Opti-MEM™ I Reduced-Serum Medium | 31985062 | Gibco™ |
| MCDB 131 Medium, no glutamine | 10372019 | Gibco™ |
| Freezing medium | 10% DMSO, 90% fetal calf serum (FCS) |  |
| Tumor cell and HEK 293T cell medium with RPMI | RPMI Medium 1640 (1x), 10% FCS |  |
| Tumor cell and HEK 293T medium DMEM/RPMI | with 10% FCS and with/ without penicillin/streptomycin |  |
| T cell (human) medium | RPMI Medium 1640 (1x), 10% FCS, 2 mM L-Glutamin |  |
| **HDRT-generation** |  |  |
| AMPure XP Beads | A63881 | Beckman Coulter |
| DpnI | R0176S | New England Biolabs |
| DynaMag2 | 12321D | Thermo Fisher Scientific |
| Kapa HiFi HotStart ReadyMix | 07958935001 | Roche |
| NEBuilder HiFi DNA Assembly Master Mix | E2621L | New England Biolabs |
| Q5 Hot Start High-Fidelity 2X Master Mix | M0494S | New England Biolabs |
| XL10-Gold Ultracompetent Cells | 200315 | Agilent |
| **mRNA** |  |  |
| mMESSAGE mMACHINE™ T7 ULTRA Transkriptionskit | AM1345 | ThermoFisher Scientific |
| mMESSAGE mMACHINE™ T7 Transkriptionskit | AM1344 | ThermoFisher Scientific |
| MEGAclear™ Kit (500µg) | AM1908 | ThermoFisher Scientific |
| Lipofectamine™ MessengerMAX™ Transfektionsreagenz | LMRNA008 | Life technologies |
| RediPlate 96 RiboGreen RNA-Kit (Sensitivity 3 - 200ng) | R32700 | ThermoFisher Scientific |
| Quant-it™ RiboGreen RNA Assay-Kit (Sensitivity 1 - 200ng) | R11490 | ThermoFisher Scientific |
| RiboRuler Low Range RNA Ladder, ready-to-use | SM1833 | ThermoFisher Scientific |
| RiboRuler High Range RNA Ladder | SM1821 | ThermoFisher Scientific |
| TheraPure™ GMP N1-Methylpseudo-UTP, 100 mM sodium solution | R0491SKB015 | ThermoFisher Scientific |
| RNeasy Micro Kit (50) (50k cells, 0- 10µg) | 74004 | Qiagen |
| RNeasy Mini Kit (50) (1mio cells, 0.5 - 35µg) | 74104 | Qiagen |
| RNeasy Mini Kit (250) | 74106 | Qiagen |
| QIAwave RNA Mini Kit (250) | 74536 | Qiagen |
| RNeasy Midi Kit (50) (30-100mio cells, 50 - 1000µg) | 75144 | Qiagen |
| QIAcuity OneStep Advanced Probe Kit (1ml) | 250131 | Qiagen |
| QIAcuity OneStep Advanced Probe Kit (5ml) | 250132 | Qiagen |
| Nuklease-freies Wasser (nicht DEPC-behandelt) | AM9937 | Life technologies |
| GeneArt™ CRISPR Nuclease mRNA | A29378 | ThermoFisher Scientific |
| tGFP mRNA for 1-Step Human IVT Kits | 88880 | ThermoFisher Scientific |
| CleanCap Cas9 mRNA | L-7606-100 | Tebubio (Trilink biotechnologies) |
|  | L-7606-1000 | Tebubio (Trilink biotechnologies) |
| CleanCap EGFP mRNA | L-7601-100 | Tebubio (Trilink biotechnologies) |
|  | L-7601-1000 | Tebubio (Trilink biotechnologies) |
| CleanCap Reagent AG | N-7113-1 | Tebubio (Trilink biotechnologies) |
| CleanCap Reagent AG | N-7113-5 | Tebubio (Trilink biotechnologies) |
| CleanCap Reagent AG | N-7113-10 | Tebubio (Trilink biotechnologies) |
| N1-Methylpseudouridine-5'-Triphosphate - (N-1081) | N-1081-1 | Tebubio (Trilink biotechnologies) |
| N1-Methylpseudouridine-5'-Triphosphate - (N-1081) | N-1081-5 | Tebubio (Trilink biotechnologies) |
| N1-Methylpseudouridine-5'-Triphosphate - (N-1081) | N-1081-10 | Tebubio (Trilink biotechnologies) |
| N1-Methylpseudouridine-5'-Triphosphate - (N-1081) | N-1081-100 | TriLink BioTechnologies, LLC |
| HiScribe T7 mRNA Kit with CleanCap Reagent AG (1,6µmol CleanCap AG/KIT) | E2080S | NEB |
| HiScribe™ T7 High Yield RNA Synthesis Kit | E2040S | NEB |
| HiScribe™ T7 ARCA mRNA Kit (with tailing) | E2060S | NEB |
| HiScribe™ T7 ARCA mRNA Kit (without tailing) | E2065S | NEB |
| Monarch® RNA Cleanup Kit (500 μg) | T2050S | NEB |
| Monarch® RNA Cleanup Kit (500 μg) | T2050L | NEB |
| Monarch® RNA Cleanup Kit (50 μg) | T2040S | NEB |
| Monarch® RNA Cleanup Kit (50 μg) | T2040L | NEB |
| RNA Loading Dye, (2X) | B0363S | NEB |
| Low Range ssRNA Ladder (50 - 1000bp) | N0364S | NEB |
| ssRNA Ladder (500 - 9000bp) | N0362S | NEB |
| Qubit™ RNA Extended Range (XR) 0,1-20µg assay kit | Q33223 | ThermoFisher Scientific |
| Qubit™ RNA Extended Range (XR) 0,1-20µg assay kit | Q33224 | ThermoFisher Scientific |
| Qubit™ RNA Broad Range (BR) 10-1200ng assay kit | Q10210 | ThermoFisher Scientific |
| Qubit™ RNA Broad Range (BR) 10-1200ng assay kit | Q10211 | ThermoFisher Scientific |
| Qubit ssDNA 0,2 bis 240 ng Assay-Kit | Q10212 | ThermoFisher Scientific |
| Qubit™ Assay-Röhrchen | Q32856 | ThermoFisher Scientific |
| **CRISPR knock-in** |  |  |
| Alt-R® S.p. Cas9 Nuclease V3, 500 µg | 1081059 | IDT |
| Gene-specific single-guide RNA (sgRNA) |  |  |
| Poly-L-glutaminsäure Natriumsalz | P4761-100MG | Sigma-Aldrich |
| Alt-R HDR Enhancer V2 | 10007910 | IDT |
| **4D Nucleofector** |  |  |
| 4D-Nucleofector® core unit | AAF-1003B | Lonza Bioscience |
| 4D-Nucleofector® X Unit | AAF-1003X | Lonza Bioscience |
| SF Cell Line 4D-NucleofectorTM X Kit L | V4XC-2024 | Lonza Bioscience |
| P3 Primary Cell 4D-NucleofectorTM X Kit L | V4XP-3024 | Lonza Bioscience |
| **Neon® Transfection System** |  |  |
| Neon Transfection System (pulse generator) | MPK5000 | Thermo Fisher Scientific |
| Neon 1-channel pipette station | MPK5000 | Thermo Fisher Scientific |
| Neon™ Transfection System Pipette | MPP100 | Thermo Fisher Scientific |
| Neon™ Transfection System 100 μL Kit  (Buffer R, Buffer T, Tubes and Tips) | MPK10096 | Thermo Fisher Scientific |
| **dPCR** |  |  |
| Pipet-Lite Multi Pipette L8-200XLS+ | 17013805 | Mettler-Toledo |
| Rainin Pipette Tips TR LTS 200 µL F 960A/10 | 17014963 | Mettler-Toledo |
| Eppendorf twin.tec® PCR Plate 96, semi-skirted, clean | 30128575 | Eppendorf |
| DANN LoBind® Tubes | 30108051 | Eppendorf |
| QIAcuity Digital PCR System (6-plex) |  | Qiagen |
| QIAamp DNA Mini Kit (50) | 51304 | Qiagen |
| QIAcuity Probe PCR Kit (5 ml) | 250102 | Qiagen |
| QIAcuity Nanoplate 26k 24-well (10) | 250001 | Qiagen |
| QIAcuity Nanoplate 26k 8-well (10) | 250031 | Qiagen |
| QIAcuity Nanoplate 8.5k 24-well (10) | 250011 | Qiagen |
| QIAcuity Nanoplate 8.5k 96-well (10) | 250021 | Qiagen |
| Nanoplate Seals (11) | 250099 | Qiagen |
| Nanoplate Tray | 250098 | Qiagen |
| XbaI | R0145S | New England Biolabs |
| **Incucyte** |  |  |
| Incucyte® Clearview 96-well Reservoir Plate | 4600 | Sartorius |
| Incucyte® Clearview 96-well Plate for Chemotaxis | 4582 | Sartorius |
| Recombinant Protein G | 101200 | Invitrogen (Thermo) |
| ICAM-1 Protein, Human, Recombinant (ECD, His & hFc Tag) | 10346-H03H | SinoBiological |
| Incucyte® Cytotox Dye for Counting Dead Cells: Green | 4633 | Sartorius |
| Incucyte® Nuclight Rapid Red Dye for Live-Cell Nuclear Labeling | 4717 | Sartorius |
| **CAR T cells and Cytokine effects** |  |  |
| CD8^+^ T cell isolation kit, human | 130-096-495 | Miltenyi |
| Pan T cell isolation kit, human | 130-096-536 | Miltenyi |
| Human CXCL10/IP-10 DuoSet ELISA, 15 Plate | DY266 | R&D |
| Human CXCL10 (IP-10) Recombinant Protein | 300-12 | Peprotec® |
| Human CXCL11/I-TAC DuoSet ELISA | DY672 | R&D |
| Human I-TAC (CXCL11) Recombinant Protein | 300-46 | Peprotec® |
| Human IFN-gamma Recombinant Protein, PeproTech® | 300-02-100UG | Thermo fisher |
| Human IFN-gamma ELISA Kit | DIF50C | R&D |
| Human IL-2 ELISA | 555190 | Becton Dickinson |
| pCMV-Rev2 (p13.33) |  | Takara |
| Viral Packaging PCHGP-2 (p14.36) |  | Takara |
| pCMV-G (p15) |  | Takara |
| UltraPure™ BSA (50 mg/mL) | AM2616 | Thermo Fisher Scientific |
| Effectene Transfection Reagent | 301425 | Qiagen |
| Opti-MEM™ I Reduced Serum Medium | 31985070 | Gibco |

#### Supplementary Table 7. Antibodies used for flow cytometry and IF Staining

| **Antibodies** | **Clone** | **Dilution** | **Catalogue no.** | **Provider** |
| --- | --- | --- | --- | --- |
| APC-Cy7 |  | 1:5000 |  | Thermo Fisher Scientific |
| Invitrogen, CD34 Monoclonal Antibody (QBEND/10), PE | 537860 | 1:10 | #MA1-10205 | ThermoFisher Scientific |

#### Supplementary Table 8. Software

| **Software** | **Use** | **Provider** |
| --- | --- | --- |
| CRISPR Design Tool | gRNA desing | Synthego <https://www.synthego.com/products/bioinformatics/crispr-design-tool> |
| FlowJo_v.10.6.2 | Analysis of flow cytometry data | Becton, Dickinson and Company; 2023 (FlowJo) |
| Gen5_v2.04 | ELISA assay measurement | BioTek |
| GeneGlobe | Target-specific PCR assay Analysis of CRISPR edits | Qiagen https://geneglobe.qiagen.com/ |
| GraphPad PRISM_v8 to v10 | Data analysis and graphical presentation | GraphPad |
| Incucyte 2021B  IncuCyte S3 | Analysis of live cell imaging | Essen Bioscience |
| ICE CRISPR Analysis Tool | Analysis of CRISPR edits | Synthego <https://www.synthego.com/products/bioinformatics/analysis> |
| NCBI Primer-BLAST | primer design and screening for specificity | [*https://blast.ncbi.nlm.nih.gov/Blast.cgi*](https://blast.ncbi.nlm.nih.gov/Blast.cgi)  *Ye, J., Coulouris, G., Zaretskaya, I. et al. Primer-BLAST: A tool to design target-specific primers for polymerase chain reaction. BMC Bioinformatics 13, 134 (2012). https://doi.org/10.1186/1471-2105-13-134* |
| Primer3Plus | Primer design and adjustment | Free software, Copyright (c) 2006, 2007 by Andreas Untergasser and Harm Nijveen |
| ProSort_v1.6 | FACS Sorter | Bio-Rad |
| QIAcuity Software Suite | dPCR analysis, calculation of absolute copy numbers | Qiagen |
| SnapGene | Development of cloning strategies, new Plasmids, primer design | Dotmatics, www.snapgene.com |
